# Molecular insights into the μ-opioid receptor biased signaling

**DOI:** 10.1101/2021.03.22.436421

**Authors:** Xiaojing Cong, Damien Maurel, Héléne Déméné, Ieva Vasiliauskaité-Brooks, Joanna Hagelberger, Fanny Peysson, Julie Saint-Paul, Jérôme Golebiowski, Sébastien Granier, Rémy Sounier

**Affiliations:** Institut de Génomique Fonctionnelle, CNRS UMR-5203 INSERM U1191, University of Montpellier, F-34000 Montpellier, France; Centre de Biochimie Structurale, CNRS UMR 5048-INSERM 1054- University of Montpellier, 29 rue de Navacelles, 34090 Montpellier Cedex, France; Université Côte d’Azur, CNRS, Institute of Chemistry of Nice UMR7272, F-06108 Nice, France; Department of Brain and Cognitive Sciences, Daegu Gyeongbuk Institute of Science and Technology, Daegu 711-873, South Korea

## Abstract

GPCR functional selectivity has opened new opportunities for the design of safer drugs. Ligands orchestrate GPCR signaling cascades by modulating the receptor conformational landscape. Our study provides insights into the dynamic mechanism enabling opioid ligands to preferentially activate the G protein over the β-arrestin pathways through the μ-opioid receptor (μOR). We combined functional assays in living cells, solution NMR spectroscopy and enhanced-sampling molecular dynamic simulations to identify the specific μOR conformations induced by G protein-biased agonists. In particular, we describe the dynamic and allosteric communications between the ligand-binding pocket and the receptor intracellular domains, through conserved motifs in class A GPCRs. Most strikingly, the biased agonists triggered μOR conformational changes in the intracellular loop 1 and helix 8 domains, which may impair β-arrestin binding or signaling. The findings may apply to other GPCR families and provide key molecular information that could facilitate the design of biased ligands.

## INTRODUCTION

Cell signaling relies on second messenger systems that are modulated by G protein-coupled receptors (GPCRs) in a ligand-specific manner. GPCRs are known for the complexity of their signaling pathways and conformational landscape. Ligands may preferentially activate or inhibit distinct signaling pathways, by changing the conformations of the GPCR (Weis and Kobilka, 2018). This is known as functional selectivity (or ligand bias), which provides fine regulations of GPCR functions and new drug design opportunities. Functional selectivity of the μ-opioid receptor (μOR) is among the most studied, in a global effort to develop safer analgesics. Opioid analgesics are efficacious and inexpensive, however, their severe side effects caused the ongoing opioid epidemic starting from the 1990s. A number of studies from 2005 to 2010 associated major opioid side effects with the β-arrestin signaling pathways (reviewed in (Raehal et al., 2011)), which has driven over a decade of research for G protein-biased μOR agonists. This led to the discovery of oliceridine (TRV130) (DeWire et al., 2013) and PZM21 (Manglik et al., 2016), two G protein-biased agonists showing less side effects than morphine in early studies. Oliceridine was approved by the FDA in 2020 for pain management, however, it still has typical opioid side effects such as nausea, vomiting, dizziness, headache, and constipation. There are high expectations on PZM21, which outperformed oliceridine and morphine in mice (Manglik et al., 2016). Nevertheless, recent findings argue that G-protein selectivity may improve analgesia and tolerance but not necessarily the side effects (Kliewer et al., 2019). Opioid-induced respiratory depression and constipation could be independent of β-arrestin signaling (Kliewer et al., 2020). It was suggested that the favorable therapeutic profiles of oliceridine and PZM21 are due to low efficacy rather than functional selectivity (Gillis et al., 2020; Yudin and Rohacs, 2019). Yet, G protein selectivity correlates with broader therapeutic windows, enabling better separations of the beneficial and adverse opioid effects (Schmid et al., 2017). A potential explanation for these discrepancies is the weak selectivity of these ligands compared to the complexity of the GPCR signaling network (Conibear and Kelly, 2019). The diversity of the systems used may also be a source of inconsistency. There is a pressing need of strongly biased and highly specific ligands, to probe the μOR signaling network for more insights. Finding such probes demands understanding the molecular mechanism of functional selectivity, which is intrinsic to the GPCR. However, GPCRs are not static on-off switches but complex molecular machines which operate through strictly regulated motions (Weis and Kobilka, 2018). Functional selectivity relies on the dynamic equilibrium of GPCR conformations, which is so far poorly understood.

X-ray crystallography and cryoEM have successfully captured inactive and active μOR states(Huang et al., 2015; Koehl et al., 2018; Manglik et al., 2012). Yet, they represent essentially a few snapshots of the vast landscape of GPCR conformations. The active states of ternary complexes exhibit a large outward displacement of the transmembrane helix 6 (TM6), which requires stabilization by G proteins or G protein-mimetics (reviewed in (Weis and Kobilka, 2018)). Conformational changes induced by ligand-binding, however, are very subtle and dynamic, as revealed by recent NMR studies on the β2-adrenergic receptor (β2AR) (Manglik et al., 2015; Nygaard et al., 2013) and on μOR (Okude et al., 2015; Sounier et al., 2015). Agonist binding initiate slight conformational changes in the ligand-binding domain (LBD), sufficient to trigger long-range conformational changes via the connector region (CR) until the intracellular coupling domains (ICD), in an allosteric and dynamic manner. Signaling partners (e.g. G proteins or β-arrestins) couple to the activated ICD and induce further large-scale opening of ICD to a fully-active state. This activation process is common in various GPCRs such as the leukotriene B4 receptor BLT2 (Casiraghi et al., 2016), the β1-adrenergic receptor (β1AR) (Isogai et al., 2016), the adenosine A_2A_ receptor (A_2A_R) (Clark et al., 2017; Eddy et al., 2018; Ye et al., 2016) and the muscarinic M2 receptor (M_2_R) (Xu et al., 2019). However, detailed activation dynamics, especially the mechanism of functional selectivity, are difficult to capture by X-ray crystallography or cryoEM. NMR spectroscopy has proven to be particularly suitable for monitoring subtle dynamic conformational transitions during GPCR activation (Kofuku et al., 2014; Liang et al., 2018; Shimada et al., 2019). MD on the other side has provided atomic-level detailed insights (Latorraca et al., 2017), such as long-timescale MD of β2AR deactivation and activation (Dror et al., 2011; Kohlhoff et al., 2014), as well as enhanced-sampling MD of the activation of M_2_R (Miao et al., 2013) and A_2A_R (Lovera et al., 2019). Here, we combined a thorough functional investigation of agonists, NMR spectroscopy and molecular dynamics (MD) simulations to obtain atomistic-level descriptions on μOR agonism. To this end, we have established a dual isotope labeling NMR scheme for μOR based on our previous study (Sounier et al., 2015). For the MD, we used the REST2 enhanced sampling scheme (REST2-MD, see METHOD DETAILS), which has proven efficient in monitoring GPCR activation (Cong et al., 2019; Cong et al., 2018; Cong and Golebiowski, 2018; Sena et al., 2017).

We studied five chemically distinct μOR agonists (**Figure 1A**): DAMGO, a well-characterized μOR-specific peptide (Emmerson et al., 1994; Handa et al., 1981); buprenorphine, a semi-synthetic thebaine analogue and partial agonist (Cowan et al., 1977a; Cowan et al., 1977b); BU72, a potent buprenorphine derivative (Neilan et al., 2004); as well as the aforementioned biased agonists, oliceridine and PZM21. By comparing the effects of these agonists on the μOR conformational dynamics, we provide here unprecedented insights into the molecular mechanism of μOR functional selectivity.

**Figure 1.**
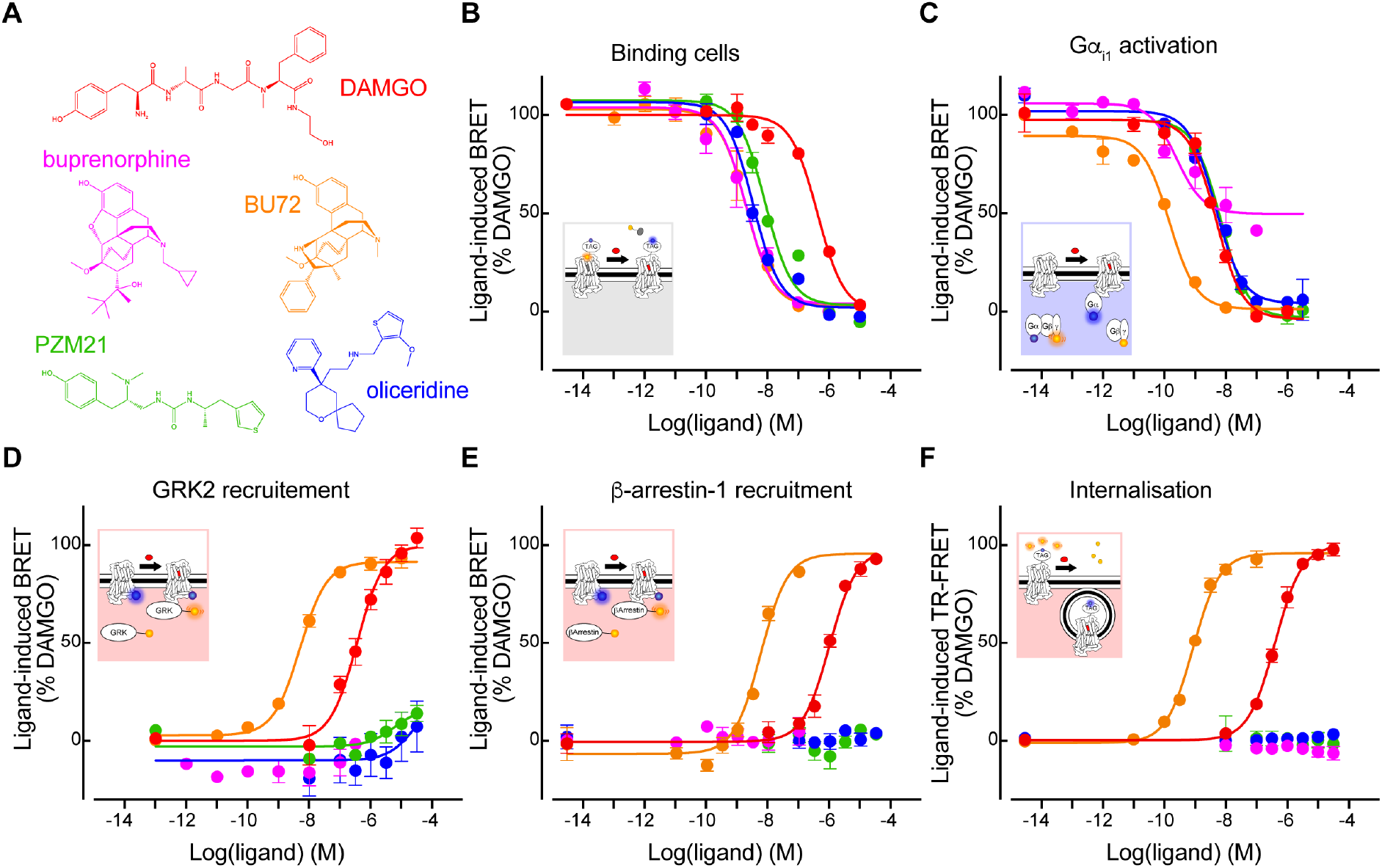
Functional characterizations of the μOR agonists. (**A**) Chemical structures of the five agonists. (**B—F**) Dose-dependent response curves of the agonists in (B) competitive binding against fluorescent naltrexone in living cells, (C) activating Gα_i1_, (D) inducing GRK2 recruitment, (E) inducing β-arrestin1 recruitment, and (F) inducing μOR internalization. Color code in (B—F) is the same than (A). Data shown are the means ± S.D. of a representative experiment performed in triplicates normalized to the maximal response induced by DAMGO and fitted using an operational model of agonism.

## RESULTS

### Opioids signaling in living cells

We first characterized the functional profiles of the five opioids in cell-based assays for their abilities to bind the target, to activate G proteins (Gi/o) and GPCR kinase 2/5 (GRK2/5), to recruit β-arrestins 1 and 2, and to trigger μOR internalization. Using advanced fluorescence methods, we probed i) competitive ligand binding to the target cells and μOR against naltrexone, ii) dissociation of the G protein heterotrimer, iii) ligand-induced μOR interactions with GRK2/5 or iv) with β-arrestins-1/2, and v) diminution of cell-surface μORs (internalization) (**Figures 1B—1F and S1; Table 1**).

All the five opioids behaved as agonists in the Gi_1-2-3_ and Go_a-b_ activation assays (**Figures 1C and S1A—S1D**). PZM21 and oliceridine behaved as full agonists in our G protein assays, similar to the data by Ehrlich *et al.* using the same assays (Ehrlich et al., 2019). However, other studies described them as partial agonists (Gillis et al., 2020; Hill et al., 2018; Yudin and Rohacs, 2019). These discrepancies highlight how functional assay outcomes may vary due to overexpression, assay conditions, assay readout amplification, or the presence of high receptor reserve (Kelly 2013), whereby an agonist may achieve maximal response by occupying only a fraction of the existing receptor population. Compared to DAMGO (our reference ligand), buprenorphine was the only partial agonist in our assays, showing 63 ± 9% to 83 ± 10 % of efficacy even at saturating concentration (**Figures 1B—1C and S1A—S1D; Table 1)**. This agrees with previous cell signaling assays (Ehrlich et al., 2019; McPherson et al., 2010; Traynor and Nahorski, 1995). Concerning the recruitments of GRK2/5 and β-arrestin-1/2, and μOR internalization, DAMGO and BU72 showed comparable efficacies with the latter being more potent, whereas oliceridine, PZM21 and buprenorphine showed nearly no response (**Figures 1D—1F and S1E; Table 1).** Therefore, oliceridine, PZM21 and buprenorphine were clearly G protein biased. Note that the bias factors could not be determined since no transduction coefficient (log(τ/*K*_A_) values could be calculated due to their lack of measurable response in the β-arrestin-1/2, GRK2/5 and internalization assays. Because our functional assays were to be compared with the NMR and REST2-MD data which provide rather qualitative information, we did not further quantify the ligands’ efficacy or bias. For the purpose of this study, we simply classified DAMGO and BU72 as unbiased full agonists, oliceridine and PZM21 as G protein-biased full agonists, and buprenorphine as a G protein-biased partial agonist.

### Development of multidomain NMR sensors for allosteric GPCR motions

We used a previously established mouse μOR construct for NMR spectroscopy, which contained an M72 T mutation (superscript refers to the Ballesteros-Weinstein numbering) that increased the expression level (Sounier et al., 2015) and reduced the NMR peak overlaps. The N-terminus was truncated before residue G52. This μOR construct maintained intact μOR functions and was stable for the duration of the NMR experiments (Sounier et al., 2015) (**Figures S1F and S2A**). Our previous approach used lysine sensors to probe μOR motions in the solvent-accessible domains (Sounier et al., 2015). Here we developed a novel dual-isotope methyl labeling scheme to monitor the solvent-accessible domains and the 7-transmembrane helices (7TM) simultaneously. For this purpose, we introduced NMR-active ^13^C probes into methionines (biosynthetically) and lysines (through reductive methylation) of the μOR construct. The heteronuclear multiple-quantum coherence (HMQC) pulse sequences were used to obtain 2D ^1^H-^13^C chemical shift correlation maps (see METHOD DETAILS). The ^13^CH_3_-ε-methionine peak assignments were obtained by mutagenesis of individual methionine residues and the dimethylamine peak assignments from our previous work (Sounier et al., 2015). We unambiguously assigned twenty sensors in the unliganded (*apo*) μOR (**Figures 2A—2C and S2-S3; see Methods**). The G52 backbone amine was at the N-terminus. Three lysines (K209^ECL2^, K233^5.39^, K303^6.58^), and four methionines (M65^1.29^, M130^2.66^, M203^4.61^, M205^4.63^) were located in the ligand binding domain (LBD). In the intracellular coupling domain (ICD), seven lysines (K98^ICL1^, K100^ICL1^, K174^ICL2^, K260^5.66^, K269^6.24^, K271^6.26^ and K344^8.51^) and four methionines (M161^3.46^, M255^5.61^, M264^ICL3^, M281^6.36^) were assigned. Only one methionine M243^5.49^ was assigned to the connector region (CR).

**Figure 2.**
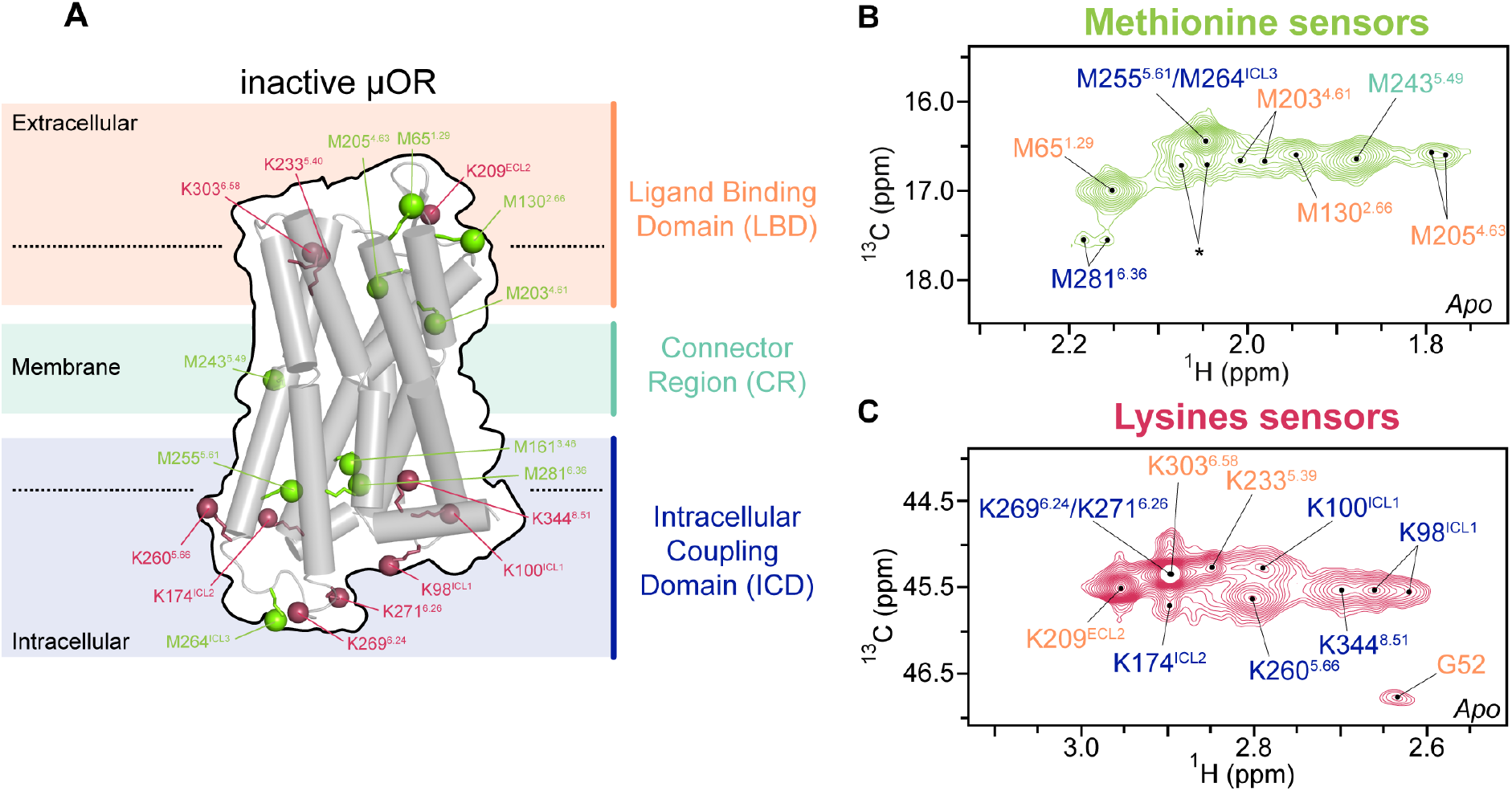
Development of multi-domain NMR sensors. (**A**) Location of the NMR sensors in a cartoon representation of μOR in inactive form. The NMR sensors, ε-CH_3_ of methionine (green) and ε-NH_2_ of lysine (raspberry), are shown in balls, in the ligand binding domain (LBD, pale orange), the connector region (CR, pale green) and the intracellular coupling domain (ICD, blue). (**B** and **C**) Extracted 2D HMQC spectra of the methionine and lysine sensors, as well as the backbone amine of the N-terminal residue G52, in *apo* μOR. Asterisk indicates the peak positions of residual resonances of the N-terminal methionine in a small amount of untruncated μOR.

To monitor agonist-induced μOR activation, we collected the HMQC spectra of μOR bound to each agonist, with and without the G protein-mimetic nanobody, Nb33 (**Figures 2A—2C and S2D—S2F**). Nb33 was necessary to reach the fully active state of μOR (Sounier et al., 2015), whereas without Nb33 we obtained the effects of the agonist binding, which more likely reflect the intrinsic mechanism of ligand bias. Taking into account the receptor concentration and the ligand depletion (Hulme and Trevethick, 2010), we estimated that 99% of the μORs should be in complexes during the NMR measurements.

### Conformational link between the ligand binding domain and the intracellular partner protein binding site

Upon binding of the agonists alone, the LBD sensors M65^1.29^ and M205^4.63^ exhibited multiple peaks with different intensities, which varied among the five agonists (**Figure S4D-S4E and S4M-S4N**). This indicates intermediate-to-slow exchanges among multiple conformations of the receptor and/or the agonists on the NMR timescales. The N-terminal sensor G52 showed a remarkable peak appearance for all the five agonists (**Figure S4G-S4H**), which was further increased upon Nb33 binding (**Figure S4I**). This suggests that ligand binding stabilized the N-terminus, which was enhanced upon Nb33 binding. Indeed, the N-terminus formed a pocket lid in the crystal structure of BU72-μOR-Nb33 (PDB 5C1M). Nb33 binding at the ICD also caused spectra changes in the extracellular loop 2 (ECL2) in a ligand-dependent manner (**Figure S4A-S4C**). These results suggest that i) Nb33 binding at the ICD has a long-range allosteric modulation of the LBD conformations; ii) Nb33 triggers cooperative interactions among the N-terminus, ECL2 and the orthosteric pocket; and iii) the agonists alone do not stabilize the fully active state even in the LBD. Importantly, these data are in line with the allosteric link between the LBD and the ICD recently reported for other class A GPCRs (Eddy et al., 2018; Liu et al., 2017; Liu et al., 2019a).

At the G protein binding site (at TM5/ICL3/TM6 in the ICD), the five agonists induced only small conformational changes (**Figures S5B, S5F and S5J)**. Nevertheless, the small changes were sufficient for the binding of Nb33, which in turn induced large conformational changes in TM5/TM6 as seen in the crystal structures of active μOR (Huang et al., 2015) (**Figures S5C, S5G and S5K)**. The magnitudes of the changes in the ternary μOR-agonist-Nb33 complexes correlated with the G-protein efficacies of the agonists (**Figure S5D and S5H**). This suggests that the capacity of an agonist to cooperate with the G proteins allosterically determines its efficacy to activate the canonical G protein signaling pathway.

### Biased, unbiased or partial agonists exhibit different binding poses

To investigate the molecular mechanism of the discrete agonist activities, we performed REST2-MD simulations (see METHOD DETAILS and **Figure S6A**) to examine how each agonist modulates the μOR conformational ensemble. All the simulations were initiated by docking the agonist to the crystal structure coordinates of μOR in an inactive form (PDB: 4DKL) (Manglik et al., 2012) without Nb33. The REST2-MD reproduced the binding poses of DAMGO and BU72 in the cryoEM/crystal structures of μOR in active forms (PDBs 6DDE (Koehl et al., 2018) and 5C1M (Huang et al., 2015), respectively, **Figure S6B**). This confirmed sufficient conformational sampling by the REST2-MD in our protocol. The biased agonists, oliceridine, PZM21 and buprenorphine, turned out to bind deeper into the CR than DAMGO and BU72. They inserted between W293^6.48^ and TM2, whereas DAMGO and BU72 bound on top of W293^6.48^ (**Figure 3A)**. The REST2-MD also captured alternative, short-lived binding poses of oliceridine and PZM21, in which the ligands bound on top of W293^6.48^ (**Figures S6C)**. These are likely transient poses in the binding process. Indeed, the HMQC spectra in the LBD indicated exchanges between different receptor/ligand conformations for all the five agonists (**Figure S4**) Whether such transient binding poses contribute to the receptor activation in the ICD is difficult to determine, since the LBD and the ICD are loosely coupled. Therefore, we focus on the ensemble changes in the receptor conformational equilibrium, without interpreting the roles of the transient binding poses.

**Figure 3.**
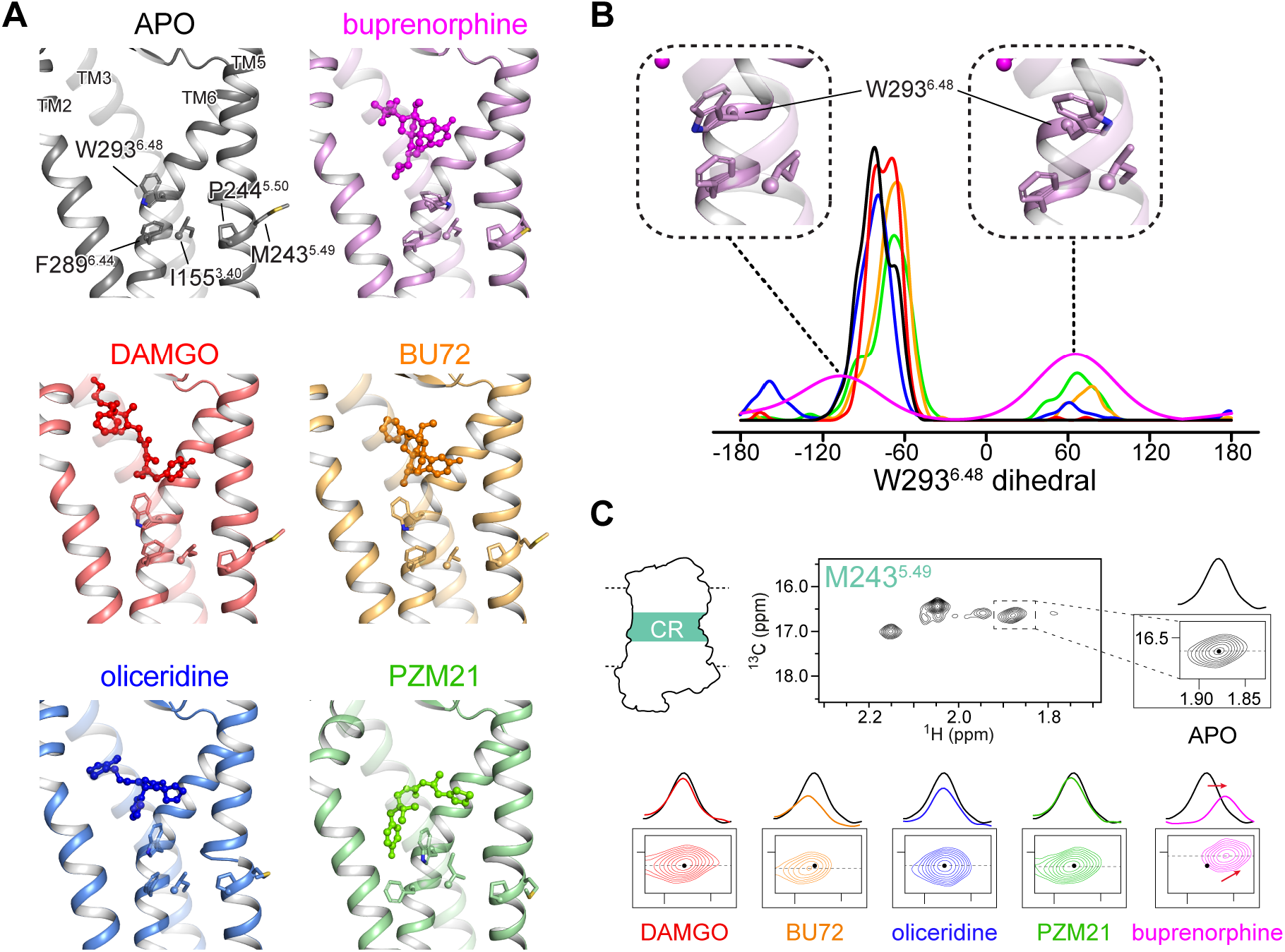
Distinct binding patterns of biased, unbiased and partial agonists in comparison with NMR. (**A**) Unbiased agonists (DAMGO and BU72) bind on top of W293^6.48^, whereas biased ones (oliceridine, PZM21 and buprenorphine) insert between W293^6.48^ and TM2. The conserved “PIF” motif (P244^5.50^, I155^3.40^ and F289^6.44^) and the NMR sensor M243^5.49^ in the CR are shown in sticks. (**B**) Ligand-dependent rotamers of W293^6.48^, as measured by the distribution of the dihedral angle *χ*_2_ during the REST2-MD simulations. (**C**) Extracted HMQC spectra of M243^5.49^ resonances in *apo* μOR (black) and at saturating concentration of DAMGO (red), BU72 (orange), oliceridine (blue), PZM21 (green) and buprenorphine (magenta). Dashed black lines indicate the position of the cross-sections shown above the spectra.

In the case of buprenorphine, the partial agonist, W293^6.48^ switched between two rotamers, whereas the full agonists maintained mostly one of the W293^6.48^ rotamers like in *apo* μOR (**Figure 3B**). W293^6.48^ is part of the conserved CW^6.48^xP motif in class A GPCRs, known as the “toggle switch” of receptor activation. It is located at the bottom of the orthosteric pocket, on top of the conserved “PIF” motif (P244^5.50^, I155^3.40^, and F289^6.44^) in the CR. Together, they form the so-called “connector region” which mediates the allosteric communications between the LBD and the ICD (Latorraca et al., 2017). W293^6.48^ plays important roles in μOR activation or inhibition (Huang et al., 2015; Yuan et al., 2015). Therefore, the different agonist binding poses with respect to W293^6.48^ may be the initial trigger of the different signaling outcomes.

The above REST2-MD observations were confirmed by the NMR sensor M243^5.49^ in the CR. We spotted a distinct pattern in the buprenorphine-μOR complex than in the other systems: the peak of M243^5.49^ shifted upfield in both the ^1^H and ^13^C dimensions, as expected from the rotation of W293^6.48^ toward M243^5.49^ upon buprenorphine binding (**Figure 3C)**. Interestingly, during an experiment using the antagonist naloxone, the M243^5.49^ peak shifted slightly in the same directions as in the case of buprenorphine (**Figure S2G**). The shift is likely associated with some constitutively active conformations in *apo* μOR that were diminished upon naloxone binding. Given the well-documented “toggle switch” role of W293^6.48^ in μOR activation (Huang et al., 2015; Yuan et al., 2015), we conclude that the low efficacy of buprenorphine is associated with its low capability to stabilize the required W293^6.48^ rotameric state for activation.

### Conformational changes in the intracellular β-arrestin binding site

The biased agonists oliceridine and PZM21 produced distinct μOR conformations in the lower half of TM7, the intracellular loop 1 (ICL1) and the helix 8 (H8) during the REST2-MD. The simulations revealed an allosteric communication from W293^6.48^ to ICL1 and H8, through the conserved motifs G^1.49^N^1.50^ and N^7.49^P^7.50^xxY^7.53^. Namely, the biased agonists inserted between W293^6.48^ and A117^2.53^ in the CR and split TM6 and TM2, which let TM7 to approach TM3 (**Figures 4A—4C**). The N^7.49^P^7.50^xxY^7.53^ motif in TM7 thus moved toward TM3, away from the G^1.49^N^1.50^ motif in TM1 (**Figures 4C—4E**). This led to remarkable inward movements of TM7 and H8 toward TM3 in the ICD, closing the cleft between H8 and ICL1 (**Figures 5A—5C**). In the case of the unbiased agonists (DAMGO and BU72), W293^6.48^ interacted with A117^2.53^ while N^7.49^P^7.50^xxY^7.53^ remained in close contact with G^1.49^N^1.50^. Residues N332^7.49^ and Y336^7.53^ formed a hydrophilic cluster with N86^1.50^ and D114^2.50^. D^2.50^ is the sodium binding site conserved in 90% of non-olfactory class A GPCRs (Katritch et al., 2014) and an important microswitch of receptor activation (Vanni et al., 2010). This configuration resembled those in the initial crystal structure (4DKL) and *apo* μOR (**Figure 4**), so did ICL1 and H8 (**Figures 5A—5C**). The partial biased agonist buprenorphine showed an in-between behavior. In the CR and the N^7.49^P^7.50^xxY^7.53^-G^1.49^N^1.50^ motifs, buprenorphine showed similar but weaker impacts than oliceridine and PZM21 (**Figures 4B and 4E**). The impacts barely reached the ICL1/H8 domains in the simulation timescale (**Figures 5A—5C**).

**Figure 4.**
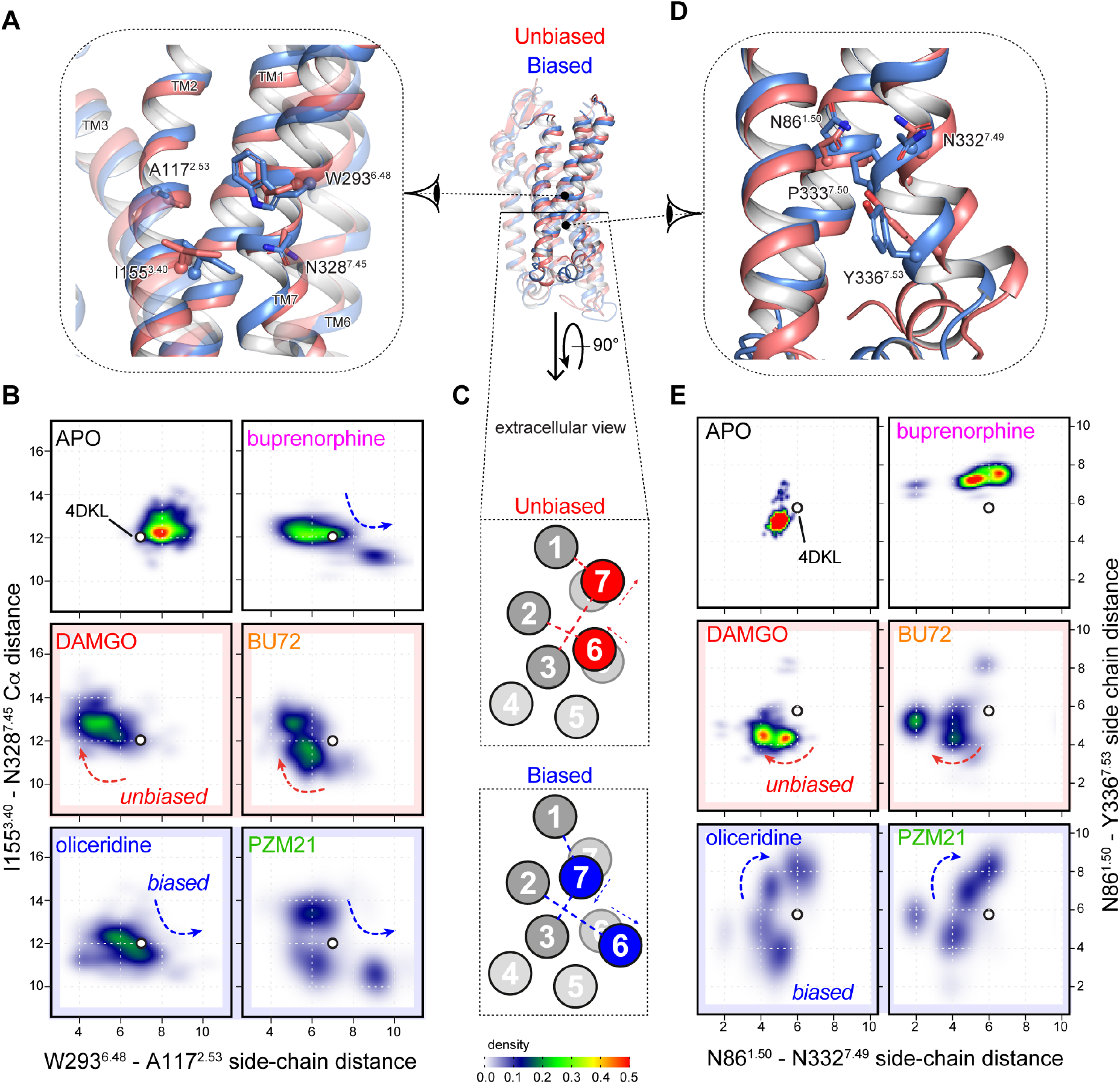
Biased agonists induced conformational changes in the CR and the lower half of TM7. (**A**) In the CR, biased agonist binding (blue) splits the side chains of W293^6.48^ and A117^2.53^, which allows TM7 to approach TM3. The movements are measured by (**B**) the minimum side-chain distances between W293^6.48^ and A1 17^2.53^, against the Cα distances between I155^3.40^ and N328^7.45^. (**C**) Schematic presentation of the inter-helical movements. Arrows indicate the direction of the movements and dashed lines indicates the distances measured for (B and E). (**D**) In the lower half of TM7, the N^7.49^P^7.50^xxY^7.53^ motif moves away from the G^1.49^N^1.50^ motif in TM1 only for biased agonists. This is measured by (**E**) the side-chain distances between N86^1.50^, N332^7.49^ and Y336^7.53^.

**Figure 5.**
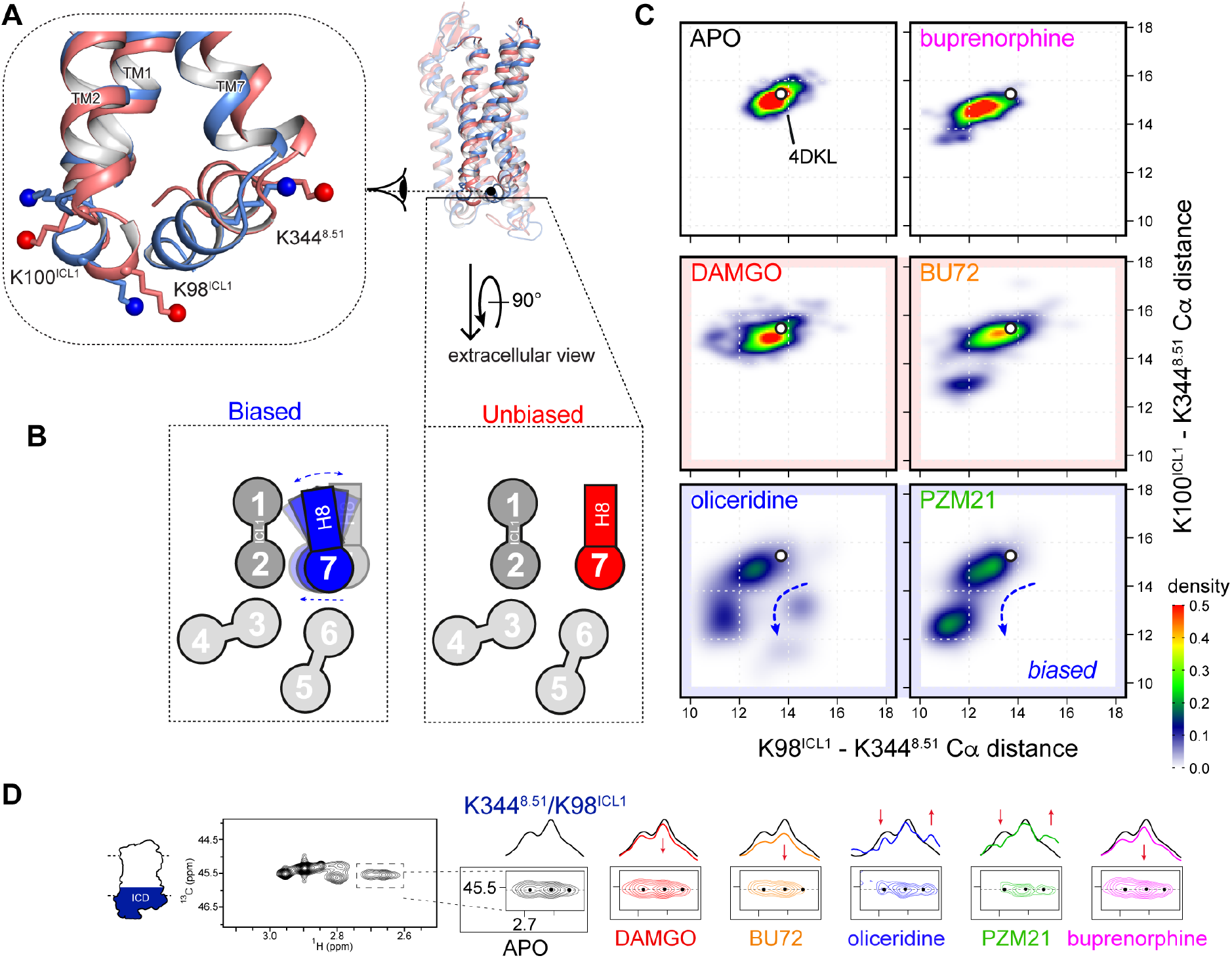
Biased agonists induced new clusters of μOR conformations in ICL1 and H8. (**A**) Biased agonist binding (blue) triggered inward movements of TM7-H8 toward TM3 and ICL1, respectively, closing the ICL1-H8 cleft. (**B**) Schematic presentation of the movements in (A) from the extracellular view, as measured by (**C**) the Cα distances between the NMR sensors K98^ICL1^, K100^ICL1^ and K344^8.51^. Density maps of the measured distances illustrate a new cluster of conformations associated with the biased agonists oliceridine and PZM21. (**D**) Extracted HMQC spectra of K344^8.51^/K98^ICL1^ resonances of μOR in *apo* form (black) and at saturating concentrations of DAMGO (red), BU72 (orange), oliceridine (blue), PZM21 (green) and buprenorphine (magenta). Dashed black lines indicate the position of the cross-sections shown above the spectra. Black dots indicate the peak centers in *apo* μOR. Red arrows indicate the changes upon agonist binding.

The NMR spectra in the ICL1/H8 domain confirmed the above findings. DAMGO, BU72 and buprenorphine binding results in the small loss of signal intensity of these sensors in H8/ICL1 (**Figure 5D**). Upon binding the biased agonists oliceridine and PZM21, the lysine sensors K98^ICL1^ and K344^8.50^ showed multiple peaks with change in signal intensity, indicating a much more complex conformational equilibrium (**Figure 5D**). This suggests that the local conformations became more dynamic. Binding of Nb33 led to signal intensity decrease for all the ligands (**Figure S5Q and S7A**). We performed REST2-MD simulations on the five receptor-agonist-Nb33 complexes and found similar patterns at ICL1 and H8 (**Figure S7B**). The distinct conformations associated with oliceridine and PZM21 were still evident in the presence of Nb33 but less remarkable. Nb33 inserts slightly between ICL1 and H8. Thus, it increased the ICL1-H8 distances in all the five complexes, while in the case of oliceridine and PZM21, it also reduced the conformational dynamics in this domain.

Overall, the results suggest that the biased agonists act on the toggle switch W293^6.48^ to trigger conformational changes in TM7, ICL1 and H8 in the ICD. This occurs in an allosteric manner, via the conserved motifs N^7.49^P^7.50^xxY^7.53^ and G^1.49^N^1.50^. The conformational changes persist even when the receptor is coupled to Nb33. Although the binding sites of G proteins and arrestins largely overlap, only arrestins interact with ICL1/H8 in the structures of arrestin in complex with rhodopsin, neurotensin receptor 1, β1AR and M2R (Huang et al., 2020; Lee et al., 2020; Staus et al., 2020; Yin et al., 2019). Therefore, the distinct TM7-ICL1-H8 conformations generated by the biased agonists likely inhibit the binding of β-arrestins but not the G proteins (Figure 6).

**Figure 6.**
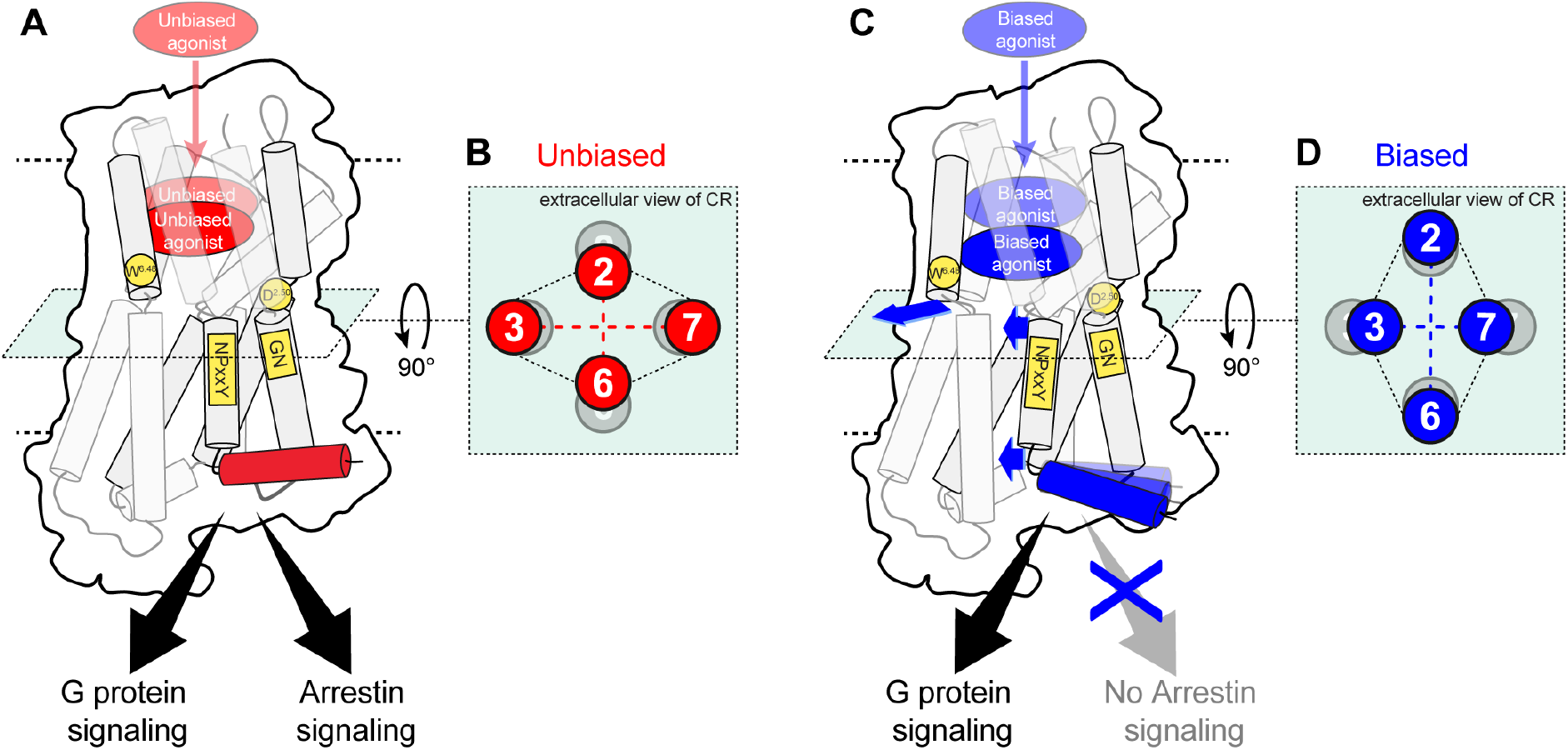
Scheme of the proposed allosteric mechanism of μOR functional selectivity. (**A**) Binding of the unbiased agonists cause W293^6.48^ in TM6 to approach TM2 in (**B**) the connector region. (**C**) Biased agonists bind deeper in the pocket and separate TM6 from TM2 in (**D**) the connector region, letting TM7 to approach TM3. This disrupts the interactions between the NPxxY motif, D114^2.50^ and the GN motif (C). The lower half of TM7 moves toward TM3, closing the cleft between H8 and ICL1, which may inhibit arrestin signaling (C). Blue arrows indicate the movements associated with allosteric functional selectivity. Conserved residues/motifs involved are highlighted in yellow. The transmembrane helices are represented by two adjoining cylinders colored in grey. The top cylinder of TM1 and TM7 are transparent for clarity.

## DISCUSSION

The pharmacological outcome of a ligand varies with the test systems and conditions, which have led to contradictory findings and the ongoing debate on whether μOR functional selectivity can separate analgesia from opioid side effects. Even the characterization of functional selectivity itself is debated, due to the lack of deep comprehensions and field standards. In this study, we provide the basic molecular mechanism of biased and partial agonism in μOR, which is intrinsic to the receptor-ligand interactions and should be consistent under different test conditions. The approach and findings can serve as high-resolution monitors for the design and evaluation of μOR ligands with specific functions, e.g. partial or biased agonism/antagonism/inverse agonism for a specific signaling pathway. Such ligands may serve to pinpoint specific aspects of the μOR signaling network, helping resolve the ongoing debates from bottom up. GPCR ligand design is challenging despite the growing number of high-resolution structures. While structure-based design and screening have lifted the hit rate, there lacks tools to design or predict biased ligands. REST2-MD can identify biased μOR agonists and thus be used for ligand screening. Although it is more costly than docking or standard MD, it provides mechanistic and dynamics insights into the ligand actions, which is essential for the subsequent ligand optimization.

This study highlights the dynamic and allosteric details of μOR pre-activation upon agonist binding, prior to the coupling of intracellular signaling partners. The pre-activation stage is crucial for drug design because it is dictated by the ligands. Nevertheless, it is highly dynamic and inaccessible to X-ray crystallography or CryoEM. We took advantage of 2D HMQC NMR and enhanced-sampling MD to capture conformational dynamic patterns during the pre-activation, which differentiate the partial and the biased agonists from the full unbiased ones. The phenomenon that biased or partial ligands stabilize distinct receptor conformations has been reported for the angiotensin II receptor 1 (during preactivation) (Wingler et al., 2019), the glucagon-like peptide-1 receptor (in fully active state) (Liang et al., 2018), the β1AR (Moukhametzianov et al., 2011; Solt et al., 2017) the A_2A_R (Huang et al., 2021; Susac et al., 2018; Ye et al., 2016) and the β2AR, either in pre-active state (biased agonism) (Liu et al., 2012) or fully active state (partial and biased agonism) (Masureel et al., 2018). Interestingly, Susac *et al.* also found different inward/outward movements of TM7 on the intracellular side upon agonist and antagonist binding, which is consistent with available A_2A_R crystal structures. However, it is unclear whether the TM7 movements of A_2A_R were associated with ligand bias. One drawback of REST2-MD is the loss of temporal information. Nevertheless, it has been shown that biased agonism in μOR is not controlled by binding or signaling kinetics, suggesting a mechanism dictated by receptor conformations (Pedersen et al., 2020)

Most of the findings here are independent of Nb33, except for the interplay observed between the agonists and Nb33 (between the LBD and the ICD), the mechanism of which remains obscure. The process of G protein binding likely determines the G protein subtype selectivity through transient GPCR-G protein interactions or intermediate conformations (Du et al., 2019; Liu et al., 2019b), but this is beyond the scope of the current study. Without Nb33 or G proteins, however, our NMR and REST2-MD results coherently and robustly illustrated the specific conformational dynamics underlying the partial and biased agonisms. The mechanism relies on highly-conserve amino-acid motifs, which may be common for other class A GPCRs.

### Limitations of the study

The current study mainly focused on the inherent mechanism of μOR ligand bias, prior to G protein or arrestin binding. One important question is how G proteins or arrestins respond to the distinct μOR conformations associated with the biased agonists. However, investigations on this aspect have been very challenging technically for both MD and NMR. The size of GPCR-agonist-G protein/arrestin ternary complexes are the major limit. Their conformational changes take place in timescales that challenge today’s all-atom MD simulations. Few high-resolution structures are available, especially for the complexes with arrestins. Such ternary complexes are also unstable under the NMR experiment conditions due to the destabilizing effect of the detergents. Therefore, studies in this aspect have been limited to G protein or arrestin surrogates (e.g. modified Gα subunits and nanobodies). The aspect of G protein/arrestin binding thus remains to be further explored.

## Supporting information

supplementary informations

## ACKNOWLEDGEMENTS

We acknowledge support from INSERM (France, S.G.), CNRS (France, H.D.), the National Institutes of Health (USA, grant NIDA-DA036246 to S.G.), the German Research Foundation (Deutsche Forschungsgemeinschaft, grant CO 1715/1-1 to X.C.) and the UCA-Jedi program (France, grant ANR-15-IDEX-01 to X.C. and J.G.). The CBS is a member of the French Infrastructure for Integrated Structural Biology (FRISBI), supported by the National Research Agency (ANR-10-INBS-05) and is a GIS-IBIsA platform. We also thank Prof. Peter Gmeiner for providing Bu72 and PZM21, and GENCI-CINES (France) for providing access to the supercomputer OCCIGEN (grant A0080711387 to J.G). All cell-based assays were carried out at the ARPEGE facility (BioCampus, Montpellier).

## AUTHOR CONTRIBUTIONS

S.G. and R.S. designed the research. J.H., I.V-B., F.P., J. S-P. and R.S. designed expressed and purified the proteins. D.M. performed and analyzed cell-based assays. H.D. and R.S. performed and analyzed NMR data. X.C. performed the MD simulations. J.G. helped X.C. analyze the MD simulation data. X.C, S.G. and R.S. wrote the manuscript. All authors contributed to editing of the manuscript.

## STAR METHODS

### KEY RESOURCES TABLE

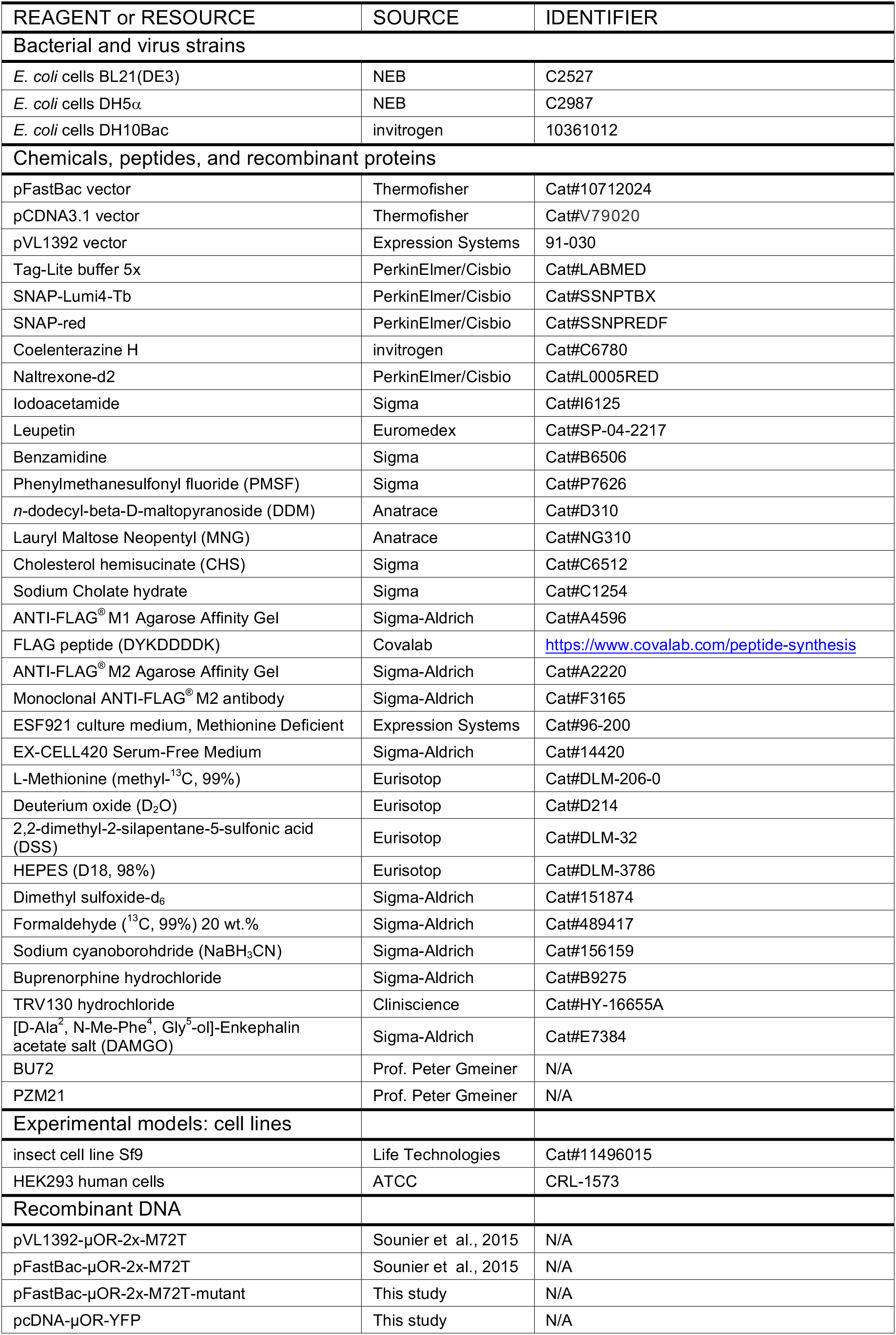
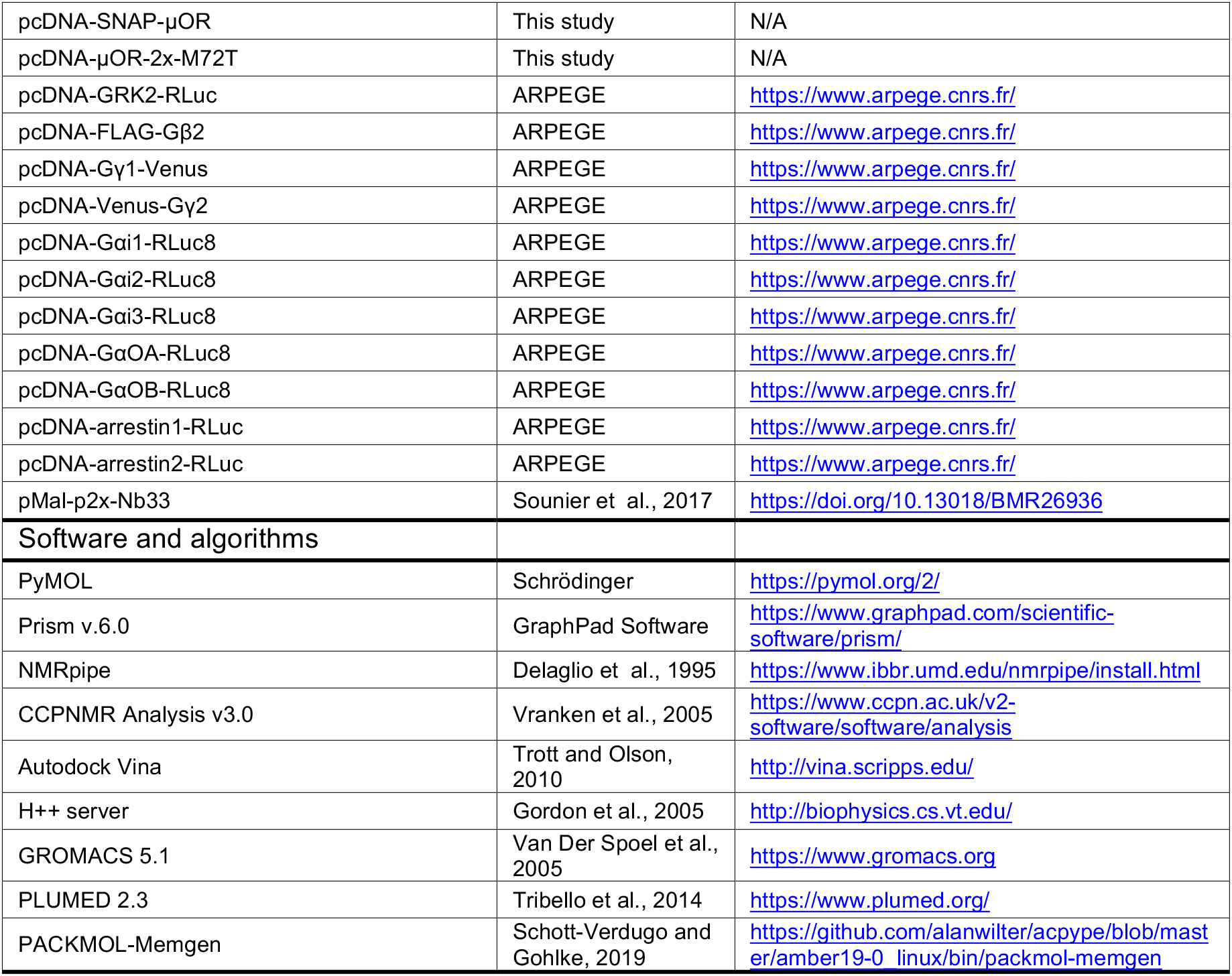

### RESOURCE AVAILABILITY

#### Lead contact

Further information and requests for reagents should be directed to and will be fulfilled by the lead contact, Rémy Sounier

#### Materials availability

DNA constructs generated by the authors can be obtained upon request from the Lead Contact, but we may require a payment and/or a completed Materials Transfer Agreement if there is potential for commercial application.

#### Data and code availability

The article includes all datasets generated or analyzed during this study.

### EXPERIMENTAL MODEL AND SUBJECT DETAILS

Mouse μOR, human Gαi1 and Gβγ were expressed in Sf9 cells infected with recombinant baculovirus (pFastBac, Invitrogen and BestBac, Expression Systems).

### QUANTIFICATION AND STATISTICAL ANALYSIS

Quantification and statistical analyses of data are described in Method Details and Figure legends.

## METHOD DETAILS

### Cell lines and transfection

HEK293 cells were grown in Dulbecco’s Modified Eagle’s Medium (DMEM, Life Technologies) supplemented with 10% fetal bovine serum (FBS, Life Technologies) without antibiotics at 37°C, 5% CO_2_. Transient transfection was performed using lipofectamine2000 (Invitrogen Life Technologies). Depending on the assay, HEK293 cells were stimulated 24h or 48h after transfection. For G protein activation and GRK2 recruitment, cells were seeded into a 6-well plate for 24h at a density of 750,000 cells per well. Cells were then detached, seeded and incubated for 24h in a 96-well white plate coated with poly-_L_-Ornithine at a density of 40,000 cells per well. For the signaling assays, we used three different constructs of full-length μOR (SNAP-μOR, μOR-2x-M72T, and μOR-YFP) depending on the experiment to be conducted (i.e. TR-FRET, BRET Gi and arrestin recruitment, respectively). For binding, β-arrestins recruitment and internalization assays, cells were directly transfected into a 96-well white plate coated with poly-_L_-Ornithine at a density of 40,000 cells per well following manufacturer’s recommended protocol. For binding assay on membrane and solubilized receptors cells were transfected using electroporation. Electroporation was performed in a volume of 400 μL with a total 5 μg SNAP-μOR plasmid and 20,000,000 cells in electroporation buffer (50 mM K_2_HPO_4_, 20 mM CH_3_COOK, and 20 mM KOH, pH 7.4). After electroporation (260 V, 1 mF, Bio–Rad Gene Pulser electroporator), cells were resuspended in 15 mL DMEM supplemented with 10% FBS in T150 culture dishes (pretreated with Poly-L-Ornithine) for 24 h.

### G protein activation assay

The cells were transfected with 4 plasmids encoding the mouse μOR receptor (Flag-μOR-2x), the β2 and Venus-γ2 G protein subunits and the Gα protein fused with a donor at a 1:1:1:1 ratio. Gαi1, Gαi2, Gαi3, GαOA and GαOB was fused to *Renilla Luciferase2* (Rluc8 provided by the ARPEGE platform). 24 h after transfection, cells were washed twice with PBS complemented with 0.9 mM CaCl_2_ and 0.5 mM MgCl_2_. Basal conditions were achieved by the addition of PBS solutions, followed by Coelenterazine H at a final concentration of 5μM. To evaluate the effects of the μOR agonists, the addition of Coelenterazine H was followed by stimulation with different agonists. Activation of the μOR promotes dissociation of the Gαβγ protein complex resulting in the Bioluminescence resonance energy transfer (BRET) signal decay. BRET between Rluc8 and Venus was measured after the addition of the Rluc8 substrate Coelenterazine H. BRET readings were collected using a Mithras LB940 plate reader (Berthold technologies, Rluc8 485 ± 20 nm; YFP 530 ± 25 nm) and the reading chamber was maintained at 37 °C throughout the entire reading time. The BRET signal was calculated by the ratio of emission of Venus (535 nm) to Rluc8 (480 nm):

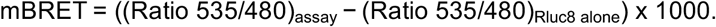

### GRK2 and GRK5 recruitment assay

The cells were transfected with 2 plasmids encoding the human μOR receptor fused to YFP at the C-terminal, and GRK2/5 was fused to *Renilla Luciferase2* (provided by the ARPEGE platform). 24 h after transfection, cells were washed twice with PBS complemented with 0.9 mM CaCl_2_ and 0.5 mM MgCl_2_. The cells were incubated with Coelenterazine H at a final concentration of 5 μM, followed by stimulation with agonists before the BRET readings were captured. Recruitment of GRK2/5 is assessed by an increase in the BRET signal.

### β-arrestins recruitment assay

The cells were transfected with 2 plasmids encoding the human μOR receptor fused to YFP at the C-terminal, and β-arrestin-1/β-arrestin-2 was fused to *Renilla Luciferase1* at the N-terminal (provided by Dr. M. Scott to the ARPEGE platform). 24 h after transfection, cells were washed twice with PBS complemented with 0.9 mM CaCl_2_ and 0.5 mM MgCl_2_. The cells were incubated with Coelenterazine H at a final concentration of 5 μM, followed by stimulation with agonists before the BRET readings were captured. Recruitments of β-arrestin-1 and β-arrestin-2 are assessed by an increase in the BRET signal.

### Internalization assay

The cells were transfected with one plasmid encoding the SNAP-μOR receptor. 24 hours after transfection, SNAP-μOR cells were washed with Tag-Lite buffer (PerkinElmer/CisBio Bioassays) and incubated at 37°C with benzylguanine-Lumi4-Tb (SNAP-Lumi4-Tb) at a concentration of 100 nM during 1 hour. The cells were washed 4 times with Tag-Lite buffer and then incubated with μOR agonists diluted in fluorescein buffer (24 μM). Reading was performed at 37°C on an Infinite F500® plate reader (TECAN, Lumi4-Terbium-criptate: 620 ± 10 nm; Fluorescein: 520 ± 10 nm) with an excitation at 337 nm and emission at 620 nm and 520 nm. Receptor internalization was monitored by time-resolved fluorescent resonance energy transfer (TR-FRET) at 37 °C during 70–80 min. The signal was calculated by the ratio of emission of terbium cryptate (620 nm) to fluorescein (520 nm): ΔR = (Ratio 620/520) X 10,000.

### Fluorescent ligand-binding assay on living cells

HEK293 cells transfected with SNAP-μOR plasmid were seeded at a density of 40,000 cells per well in 96-well white plates coated with poly-_L_-Ornithine. 24h after transfection, SNAP-μOR was labeled 1 h at 37°C with 100nM SNAP-Lumi4-Tb diluted in Tag-lite labeling buffer. Fluorescent naltrexone and agonists were diluted in Tag-lite labeling buffer. A fixed concentration of fluorescent naltrexone-d2 was determined (0.5 nM = K_d_) and used. Increasing concentration of agonists was added prior to the addition of fluorescent naltrexone-d2 in the plates containing labeled cells. Plates were incubated overnight at 4°C before homogenous time-resolved Fluorescent (HTRF) signal detection. HTRF detection was performed on a PHERAstar (BMG labtechnologies). The signal was collected both at 665 nm and 620 nm. HTRF ratio was obtained by dividing the acceptor signal at 665 nm by the donor signal at 620 nm and multiplying this value by 10,000. Data obtained were then analyzed using GraphPad Prism (GraphPad Software, Inc., San Diego, CA).

### Membrane preparation and SNAP labeling

Twenty-four hours after transfection cells were washed once with PBS solution, scraped and then collected by centrifugation 5 minutes at 300 g. The cell pellet was resuspended in 20 mL lysis buffer (10 mM HEPES, 1 mM EDTA) and homogenized using an electric homogenizer on ice. After centrifugation 5 min at 1,000g at 4 °C, the pellet was discarded and the supernatant was centrifuged at 30,000 g for 30 min at 4 °C. The resulting pellet was resuspended in 2 mL of Tag-Lite buffer containing 300 nM BG-Lumi4-Tb and incubated for 1 hour at 4 °C under circle rotator. Membranes were then resuspensed in 2 mL PBS to remove the excess of BG-Lumi4-Tb and centrifuged at 30,000 g for 30 min on a bentchtop centrifuge. This step was reproduced twice. Protein concentration was determined by BCA using BSA as standard. SNAP-μOR membranes were aliquoted and stored at −80 °C.

### Fluorescent ligand-binding assay on solubilized receptors

SNAP-μOR membranes were resuspended in solubilization buffer (see below for composition details) and stirred 1 hour at 4 °C. After centrifugation at 36,000 g for 20 min at 4 °C, the solute material was complemented with 2 mM CaCl_2_ and loaded on M1 antibody Resin. Detergent exchange protocol was performed (see below for details) and solubilized SNAP-μOR was eluted from M1 antibody resin. Naltrexone-d2 binding affinity towards solubilized SNAP-μOR was determined at 3 nM against 0.5 nM on living cells and 1 nM HEK293 cell membranes. Freshly prepared solubilized SNAP-μOR were incubated with increasing concentrations of agonists and a fixed concentration of fluorescent naltrexone-d2 (12 nM = K_d_) before HTRF signal detection.

### (Met-ε)-[^13^CH_3_]-μOR-2x M72T expression

We generated a μ-OR mouse construct with features designed to enhance stability for NMR spectroscopy. A tobacco etch virus (TEV) protease recognition site was introduced after residue 51, and a human rhinovirus 3C protease site after residue 358. Receptor expression was largely improved by using a M72^1.36^T single-point mutation as previously reported (Sounier et al., 2015). A FLAG tag was added to the amino terminus and an 8 × His tag was appended to the carboxy terminus (μOR-2x). Recombinant baculoviruses were generated using the BestBac baculovirus system according to manufacturer’s instructions (Expression Systems). For the assignment of all other mutants, we used the pFastBac baculovirus system (ThermoFischer). High titer baculoviruses encoding μOR-2x genes were used to infect *Sf9* cells at a cell density of 4 × 10^6^ cells per ml in suspension in methionine deficient media (Expression System) in the presence of 3 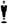 M naloxone with ^13^C methyl labelled methionine (Cambridge Isotope) added into the media at 250 mg·L^-1^ concentration. Cells were harvested by centrifugation 48 h post-infection and stored at −80 °C until purification.

### (Met-ε)-[^13^CH_3_]-μOR-2x M72T purification

Cell pellets were resuspended in 10 mM Tris-HCl pH 7.5, 1 mM EDTA buffer containing 2 mg·mL^-1^ iodoacetamide and protease inhibitors without salt to lyse the cells by hypotonic lysis. Lysed cells were centrifuged (38,420 g) and the membranes were solubilized durinf 1h @ 4 °C using buffer containing 20 mM HEPES (pH 7.5), 200 mM NaCl, 0.5% (w/v) n-dodecyl-β-D-maltoside (DDM, Anatrace), 0.3% (w/v) CHAPS, 0.03% (w/v) choslesteryl-hemi-succinate (CHS, Sigma), 2 mg·mL^-1^ iodoacetamide and protease inhibitors. The solubilized receptor was loaded onto anti-Flag M1 column (Sigma) and washed thoroughly with DDM buffer containing 20 mM HEPES (pH 7.5), 100 mM NaCl, 0.1% (w/v) DDM, 0.03% (w/v) CHAPS, 0.015% (w/v) CHS and 2 mM CaCl_2_. While on the M1 antibody resin, the receptor was exchanged into lauryl-maltose-neopentyl-glycol (MNG-14, Anatrace) detergent-containing buffer composed of 20 mM HEPES (pH 7.5), 100 mM NaCl, 0.5% (w/v) MNG-14 and 0.01 % CHS. The detergent exchange was performed by washing the column with a series of seven buffers (3 CV each) made up of the following ratios (v/v) of MNG-14 buffer and DDM buffer: 0:1, 1:1, 4:1, 9:1, 19:1, 99:1 and MNG-14 exchange buffer alone. The column was then washed with 20x critical micelle concentration (cmc) MNG-14 buffer containing 20 mM HEPES (pH 7.4), 100 mM NaCl, 0.02% (w/v) MNG and 0.0004 % CHS and the bound receptor was eluted in the same buffer supplemented with 0.2 mg·mL^-1^ Flag peptide. To remove flexible amino and carboxy termini, TEV and 3C protease were added at a 1:5 and 1:10 protease:μOR-2x ratio by weight. The sample was incubated at 4 °C overnight in the presence of 100 μM of TCEP. We then used a negative Ni-NTA chromatography step to remove TEV and 3C proteases.

### (Met-ε)-[^13^CH_3_]-μOR-2x M72T assignment procedure

To obtain sequence-specific assignments, we introduced single point mutations of 13 methionines in μOR by site-directed mutagenesis (Genecust). To identify suitable methionine substitutions, we performed a sequence alignment of μOR homologs and selected the most common amino acid for each position: M65T, M72T, M90I, M99L, M130L, M151I, M161I, M203I, M205L M243V, M255I, M264L and M281L. The mutants were expressed and purified as described above, except for M151I which was unstable in the detergent micelles. Ten methionines were unambiguously assigned by comparing the spectra of μOR and the μOR mutants in both *apo* and fully active states (Figure S3). M90 and M99 could not be assigned (Figure S3).

### (Met-ε)-[^13^CH_3_]-μOR-2x M72T reductive methylation

Receptor preparation from the Ni-NTA flow through were incubated at 4 °C overnight with 10 mM ^13^C-formaldehyde and 10 mM NaBH_3_CN. Excess of reagent was eliminated by dialysis and (Met-ε)-[^13^CH_3_],(Lys-Nε,Nε)[^13^CH_3_,^13^CH_3_]-μOR ((^13^C-M^e^,K^me2^)-μOR) was further purified by SEC chromatography in a buffer containing 0.01% MNG, 0.001% CHS, 20 mM HEPES pH 7.4 and 40 mM NaCl. The monodisperse peak was then concentrated to 30 to 60 μM final, and dialysed in 98.85% D2O buffer with 0.01% MNG, 0.0004% CHS, 20 mM HEPES-d18 pH 7.4 (uncorrected) and 40 mM NaCl.

### Nanobody Nb33 expression and purification

The Nb33 was expressed and purified as described in our previous work (Sounier et al., 2017). Briefly, The DNA sequence of Nb33 was subcloned into a pMalp2x vector containing an N-terminal, 3C protease-cleavable maltose binding protein (MBP) tag and a C-terminal 8 × His tag. Plasmids were transformed into BL21(DE3) cells and protein expression induced in liquid broth (LB) by addition of IPTG to 0.5 mM at an OD_600_ of 0.6. Cells were harvested after overnight growth at 20°C by centrifugation at 6,000 g for 30 min. Cells were resuspended in 20 mM HEPES buffer (pH 7.5), 500 mM NaCl, 0.1 mg·mL^-1^ lysozyme and PMSF was added as a protease inhibitor before lysis by sonication. The cell lysate was centrifuged at 38,420 g for 30 min at 4 °C. The soluble fraction was isolated and was supplemented with imidazole to a final concentration of 20 mM. MBP–nanobody fusions were purified by Ni-NTA chromatography and MBP was removed using 3C protease. Cleaved MBP was separated from the nanobody by additional amylose purification and size exclusion chromatography in a buffer containing 20 mM HEPES pH 7.4 and 0.1 M NaCl.

### NMR Spectroscopy

Final samples (~270 μl at 30–60 μM) were loaded into Shigemi microtubes susceptibility matched to D_2_O. All data for ligands and mutant studies were acquired on 700 MHz Bruker Avance III spectrometers (Bruker, Rheinstetten, Germany), equipped with 5 mm cryogenic H/C/N/D probes with *z* axis gradient. ^1^H-^13^C correlation spectra were recorded using heteronuclear multiple-quantum coherence (HMQC) experiments in echo anti-echo mode. ^13^C and ^1^H chemical shifts and peak line widths in the HMQC spectra reveal the chemical and magnetic environments of the ^13^C-methyl probes as well as their dynamic properties. Spectral widths in ω1 and ω2 were 8,417.5 Hz and 3,518.6 Hz at 700 MHz centred at 40 p.p.m. or 20 p.p.m. in the ^13^C dimension. ^13^C decoupling was performed with a GARP4 sequence. Typically, 134 complex points with 32–48 scans per FID were recorded, to ensure a 27-Hz resolution per point at 700 MHz before zero filling. The relaxation delay was set to 1.5 s. Thirty-two steady-state scans preceded data acquisition. Total collection time varied between 3 and 4 h, depending on the sample concentration. All the data were processed in the same manner using NMRPipe/NMRDraw (Delaglio et al., 1995). Prior to Fourier transformation, the data matrices were zero-filled to 1024 (*t*_1_) x 4096 (*t*_2_) complex points and multiplied by a sine-bell window function in each dimension. Peak fitting analysis was performed with the program nlinLS (part of the NMRDraw package) using the same approach as previously described (Okude et al., 2015; Sounier et al., 2015). The peak intensities were normalized to the volume difference between the *apo* state and the ternary complex condition (DAMGO-Nb33) as the 100%. The spectra were visualized using CCPNMR (Vranken et al., 2005). All ligands were dissolved in perdeuterated dimethyl d_6_-sulfoxide (d_6_-DMSO, Cambridge Isotope) to 100 mM and directly added to the sample in the Shigemi tube at a final concentration five-fold of receptor. Nb33 were concentrated to 0.6 mM and dialysed in 100% D_2_O buffer with 0.01% MNG, 0.001% CHS, 20 mM HEPES-d18 pH 7.4 (uncorrected) and 40 mM NaCl. The nanobodies were added directly in the Shigemi tubes at a final concentration of two-fold of the receptor before data acquisition.

### Molecular dynamics simulations

The initial coordinates of μOR were from the crystal structure of an inactive form (PDB: 4DKL). The ligands were docked to the initial μOR structure using Autodock Vina (Trott and Olson, 2010). Residues in the putative ligand-binding pocket were set flexible during docking. The protonation state of titrable residues were predicted at pH 7.4 using the H++ server (Gordon et al., 2005). The receptor-odorant complexes were embedded in a bilayer of POPC using PACKMOL-Memgen (Schott-Verdugo and Gohlke, 2019). Each system was solvated in a periodic 75 × 75 × 105 Å^3^ box of explicit water and neutralized with 0.15 M of Na^+^ and Cl^-^ ions. Effective point charges of the ligands were obtained by RESP fitting (Wang et al., 2000) of the electrostatic potentials calculated with the HF/6-31G* basis set using Gaussian 09 (Frisch et al., 2009). The Amber 99SB-ildn (Lindorff-Larsen et al., 2010), lipid 14 (Dickson et al., 2014) and GAFF (Wang et al., 2004) force fields were used for the proteins, the lipids and the ligands, respectively. The TIP3P (Jorgensen et al., 1983) and the Joung-Cheatham (Joung and Cheatham, 2008) models were used for the water and the ions, respectively.

After energy minimization, all-atom MD simulations were carried out using Gromacs 5.1 (Van Der Spoel et al., 2005) patched with the PLUMED 2.3 plugin (Tribello et al., 2014). Each system was gradually heated to 310 K and pre-equilibrated during 10 ns of brute-force MD in the *NPT-ensemble.* The replica exchange with solute scaling (REST2) (Wang et al., 2011) technique was employed to enhance the sampling with 48 replicas in the *NVT* ensemble. REST2 is a type of Hamiltonian replica exchange simulation scheme, which performs many replicas of the same MD simulation system simultaneously. The replicas have modified free energy surfaces, in which the barriers are easier to cross than in the original system (**Figure S6A**). By frequently swapping the replicas during the MD, the simulations “travel” on different free energy surfaces and easily visit different conformational zones. Finally, only the samples on the original free energy surface are collected. The replicas are artificial and are only used to overcome the energy barriers. REST2, in particular, modifies the free energy surfaces by scaling (reducing) the force constants of the “solute” molecules in the simulation system. The protein and the ligands were considered as “solute”–the force constants of their van der Waals, electrostatic and dihedral terms were subject to scaling–in order to facilitate their conformational changes. The effective temperatures used here for generating the REST2 scaling factors ranged from 310 K to 700 K, following a distribution calculated with the Patriksson-van der Spoel approach (Patriksson and van der Spoel, 2008). Exchange between replicas was attempted every 1000 simulation steps. This setup resulted in an average exchange probability of ~40%. We performed 50 ns × 48 replicas of MD in the *NVT* ensemble for each system. The first 20 ns were discarded for equilibration. From our past experiences on REST2-MD of GPCR conformational changes (Cong et al., 2019; Cong et al., 2018; Cong and Golebiowski, 2018; Sena et al., 2017), we estimated that 50 ns should achieve millisecond timescale sampling. The original unscaled replica (at 310 K effective temperature) was collected and analyzed. Cluster analysis of the ligand binding pose was carried out on the non-restrained trajectory using the Gromacs Cluster tool. The middle structure of the most populated cluster was selected as the final binding pose.

