## supplementary informations for "Molecular insights into the μ-opioid receptor biased signaling"

### Supplemental Information

| Emax (% DMAGO) | DAMGO | BU72 | Oliceridine | PZM21 | Buprenorphine |
|---|---|---|---|---|---|
| **Gαi1** | 100 | 127 ± 17 | 94 ± 5 | 95 ± 6 | 63 ± 9 |
| **Gαi2** | 100 | 109 ± 8 | 98 ± 4 | 94 ± 5 | 66 ± 18 |
| **Gαi3** | 100 | 114 ± 14 | 94 ± 3 | 100 ± 5 | 83 ± 10 |
| **GαoA** | 100 | 120 ± 17 | 87 ± 5 | 90 ± 5 | 70 ± 5 |
| **GαoB** | 100 | 120 ± 14 | 91 ± 8 | 93 ± 9 | 67 ± 6 |
| **GRK2** | 100 | 98 ± 19 | N.D. | N.D. | N.D. |
| **GRK5** | 100 | 92 ± 8 | N.D. | N.D. | N.D. |
| **β-arrestin-1** | 100 | 98 ± 10 | N.D. | N.D. | N.D. |
| **β-arrestin-2** | 100 | 96 ± 12 | N.D. | N.D. | N.D. |
| **Internalization** | 100 | 96 ± 5 | N.D. | N.D. | N.D. |
| **$pEC_{50}$** | | | | | |
| **Gαi1** | 8.11 ± 0.13 | 9.76 ± 0.14 | 8.22 ± 0.09 | 8.22 ± 0.23 | 9.29 ± 0.77 |
| **Gαi2** | 8.33 ± 0.26 | 9.92 ± 0.24 | 8.59 ± 0.35 | 8.49 ± 0.33 | 8.90 ± 0.57 |
| **Gαi3** | 8.33 ± 0.22 | 9.99 ± 0.17 | 8.44 ± 0.05 | 8.51 ± 0.23 | 8.90 ± 0.51 |
| **GαoA** | 8.14 ± 0.20 | 9.55 ± 0.25 | 8.10 ± 0.09 | 8.12 ± 0.12 | 9.02 ± 0.72 |
| **GαoB** | 8.17 ± 0.24 | 10.10 ± 0.66 | 8.27 ± 0.32 | 8.25 ± 0.24 | 8.57 ± 0.21 |
| **GRK2** | 6.16 ± 0.22 | 8.19 ± 0.16 | N.D. | N.D. | N.D. |
| **GRK5** | 6.08 ± 0.13 | 7.65 ± 0.21 | N.D. | N.D. | N.D. |
| **β-arrestin-1** | 5.05 ± 0.17 | 8.40 ± 0.32 | N.D. | N.D. | N.D. |
| **β-arrestin-2** | 6.32 ± 0.12 | 9.00 ± 1.06 | N.D. | N.D. | N.D. |
| **Internalization** | 6.46 ± 0.17 | 8.96 ± 0.16 | N.D. | N.D. | N.D. |
| **$K_i$ (nM)** | | | | | |
| **cells** | 129 ± 50 | 0.7 ± 0.3 | 3.1 ± 2.5 | 6.7 ± 5.4 | 0.3 ± 0.2 |
| **Solubilized μOR** | 1255 ± 221 | 0.9 ± 0.7 | 66.0 ± 44.0 | 59.0 ± 33.0 | 2.3 ± 2.4 |

**Table 1. Potency, $E_{max}$ (relative to the maximal response of DAMGO) and affinity of the five ligands in assays. Related to Figures 1 and S1.**

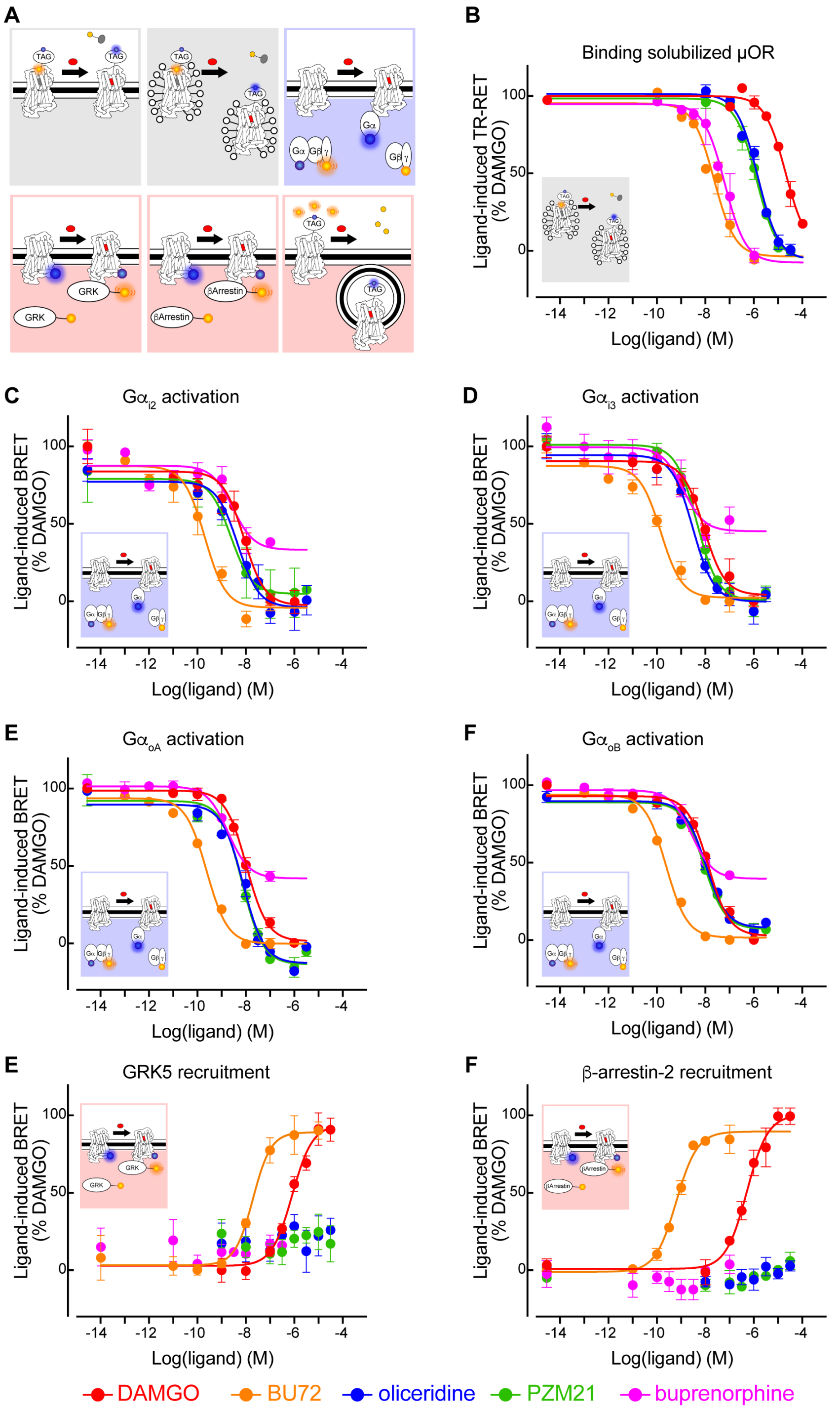


**Figure S1. Functional characterization of the five ligands. Related to Figure 1.**

(**A**) Schematic representation of the different BRET and TR-FRET assays used (donors are in blue and acceptors are in yellow). Dose-dependent response curves of the agonists in (**B**) competitive binding to solubilized μOR against fluorescent naltrexone, (**C**) activating $G\alpha_{i2}$, (**D**) activating $G\alpha_{i3}$, (**E**) activating $G\alpha_{oA}$, (**F**) activating $G\alpha_{oB}$, (**E**) inducing GRK5 recruitment, and (**F**) inducing β-arrestin-2 recruitment. Data shown are the means ± S.D. of a representative experiment performed in triplicates normalized to the maximal response induced by DAMGO and fitted using an operational model of agonism.

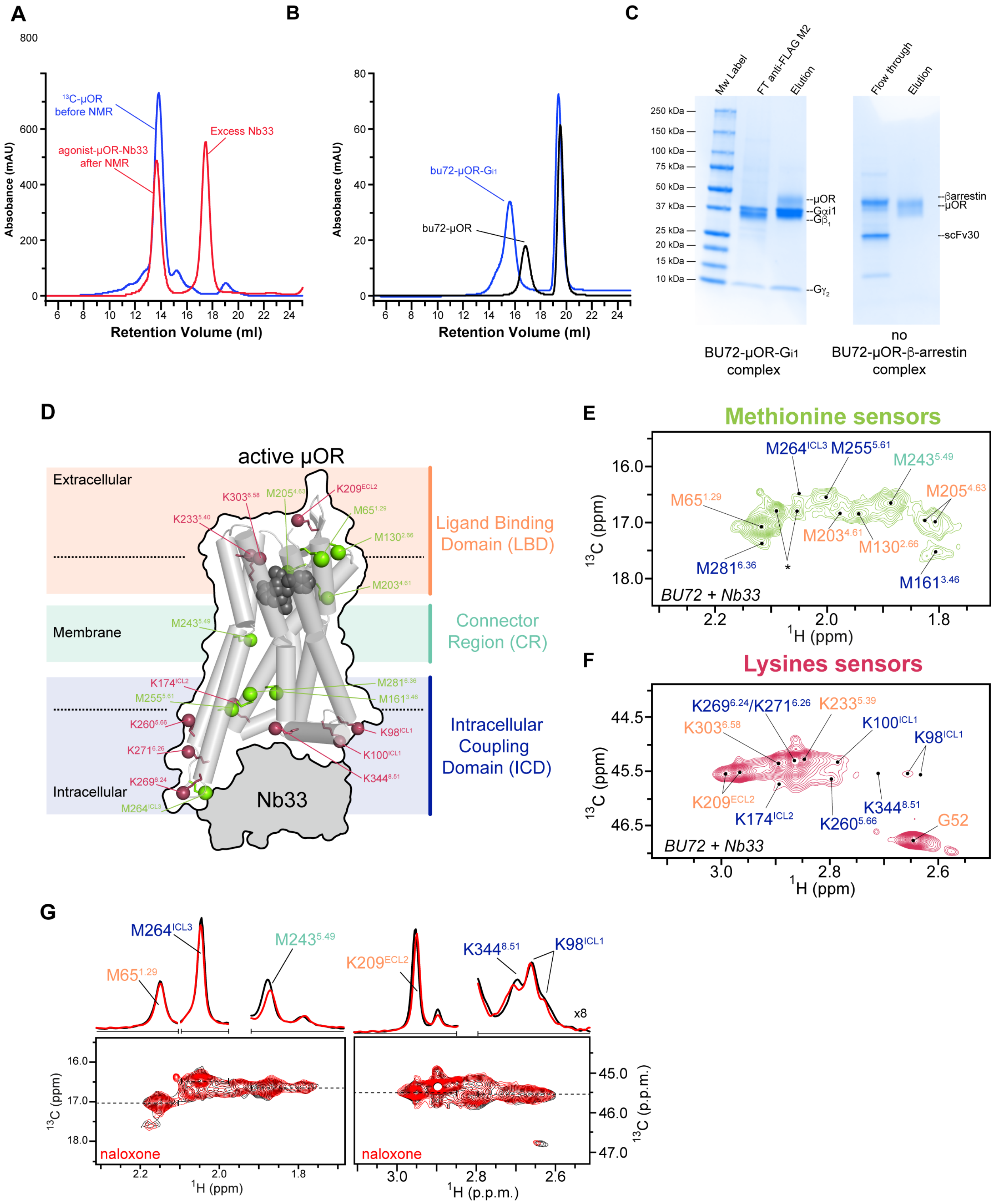


**Figure S2. Biophysical characterization of μOR complexes and development of NMR sensors. Related to Figure 2.**

(**A**) Typical size-exclusion chromatography (SEC) results showed the monodispersity of $^{13}$C-labelled μOR ($^{13}$C-μOR) before NMR (Blue) and the agonist-μOR-Nb33 complex after NMR experiments (Red) using Superdex200 columns. (**B**) Typical SEC results showed the BU72-μOR-Gi complex using Superose6 columns. (**C**) SDS-PAGE of BU72-μOR-Gi1 complexes after anti-FLAG M2 resin pull-down confirmed the complex is made of μOR, Gαi1, Gβ1, Gγ2. In comparison, BU72-μOR-arrestin complexes were not observed under the same condition. (**D**) Location of the NMR sensors in a cartoon representation of the BU72-μOR-Nb33 ternary complex. NMR sensors, ε-$CH_3$ of methionine (green) and ε-$NH_2$ of lysine (raspberry), are shown in balls. (**E** and **F**) Extracted 2D HMQC spectra of the methionine and lysine sensors, as well as the backbone amine of the N-terminal residue G52, in the BU72-μOR-Nb33 ternary complex. Asterisk indicates the peak positions of residual resonances of the N-terminal methionine in a small amount of untruncated $^{13}$C-μOR. (**G**) Comparison of HMQC spectra of μOR in apo-state (black) and saturating concentration of naloxone (red). 1D slices of HMQC spectra in the $^{1}$H dimension is shown on top of each spectra.

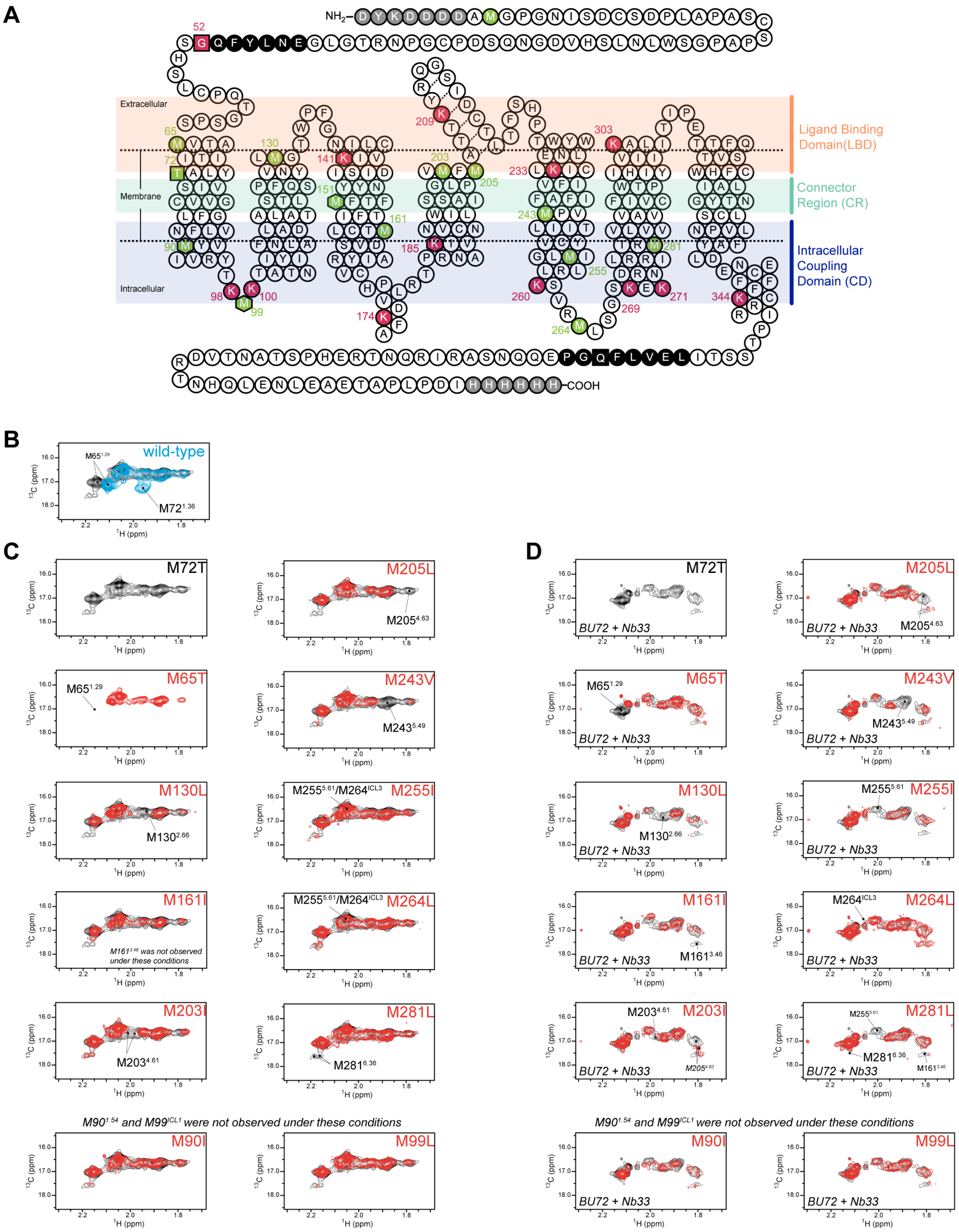
A
NH2
COOH
Extracellular
Membrane
Intracellular
Ligand Binding Domain(LBD)
Connector Region (CR)
Intracellular Coupling Domain (CD)
B
wild-type
M65^1.29
M72^1.36
C
M72T
M65T
M130L
M161I
M161^3.46 was not observed under these conditions
M203I
M205L
M243V
M255I
M264L
M281L
M90^1.54 and M99^ICL1 were not observed under these conditions
M90I
M99L
D
BU72 + Nb33
M90^1.54 and M99^ICL1 were not observed under these conditions
13C (ppm)
1H (ppm)

**Figure S3. Assignments of resonances from methyl methionines of μOR in *apo* state and in complex with BU72 and Nb33. Related to STAR Methods.**

(**A**) Snake plot of μOR sequence showing the FLAG and 6x Histidine tags (light gray), the protease cleavable motifs (black), the M72$^{1.36}$T mutation site (green, squared), as well as the methionine (green), lysine (raspberry) and G52 amine (red, squared) sensors. Unassigned residues in the 2D HMQC spectra are shown in hexagonal. (**B**) Comparison of the HMQC spectra of $^{13}CH_3$-ε-methionine labeled wild-type μOR (blue) and μOR-M72T (black) in *apo* state. (**C** and **D**) Spectra of methionine mutants in *apo* state and in the BU72-μOR-Nb33 ternary complex. The spectra of μOR-M72T (black) was used as reference to superimpose those of the other mutants (red) We highlighted the peak disappearance for each mutant. We generated and recorded the NMR spectra for 12 mutants of 13 endogenous methionines (labeled in A). Only M151I could not be recorded because of instability issues. The spectra of M90I and M99L were not assigned under the conditions used.

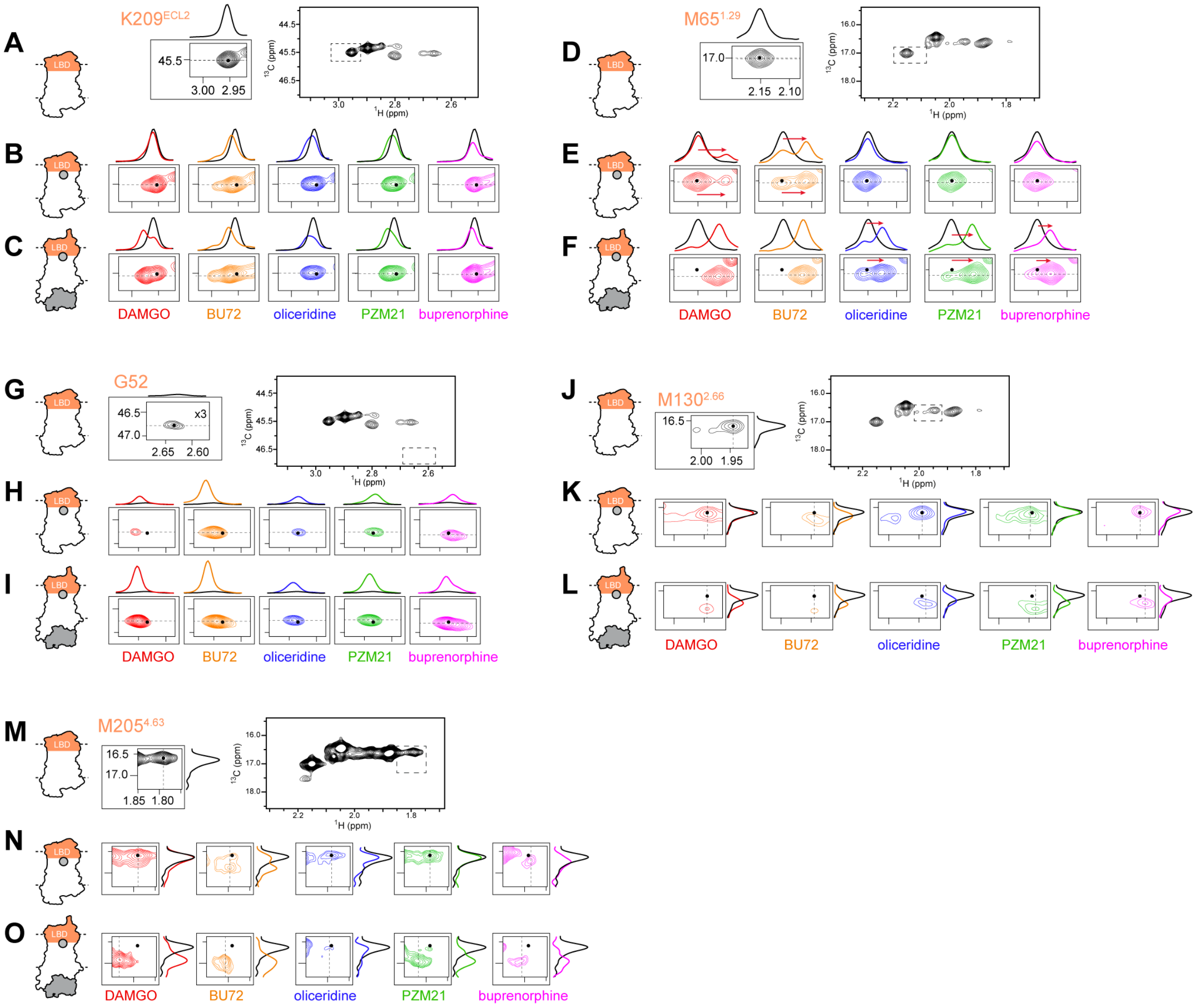


**Figure S4. NMR spectral changes in the ligand-binding domain upon binding with each agonist and Nb33 at saturating concentrations. Related to Figure 3.**

Extracted HMQC spectra of the sensors (**A—C**) K209$^{ECL2}$, (**D—F**) M65$^{1.29}$, (**G—I**) G52 amine at the truncated N-terminus), (**J—L**) M130$^{2.66}$, and (**M—O**) M205$^{4.63}$ are compared with *apo* μOR (black) (top panel), either bound to agonist (middle panel) or to agonist and Nb33 (bottom panel). Dashed black lines indicate the position of the cross-sections shown on top or on the right of the spectra.

The K209$^{ECL2}$ (**A-B**) and M130$^{2.66}$ (**J-K**) probes only sensed slight effects of ligands alone with a decrease of signals intensity and a downfield shift of the chemical shift in the $^{1}$H dimension and $^{13}$C dimension. By opposition, for those two sites, Nb33 binding to the agonist-bound receptor induced important changes both in intensity and $^{1}$H chemical shift for all the tested ligands (**C** and **L**). The N-terminus G52 probe sensed the binding of all ligands with a peak appearance (2.64 ppm and 46.75 ppm in $^{1}$H and $^{13}$C dimension) further increased in intensity upon Nb33 binding (**G-I**). These results suggest that the N-terminus of the apo form of the receptor sample more conformations than bound states. Of note, Nb33 binding at the ICD further increased the G52 methyl peak, suggesting a further stabilization of the N-terminus. Interestingly, the M65$^{1.29}$ probe revealed a ligand-specific regulation in which only BU72 and DAMGO by themselves were able to modify the chemical shift obtained in ligand free preparation (**D-F**). In particular, BU72 and DAMGO promoted the appearance of a new peak at 2.12 ppm in the $^{1}$H dimension **(E)**. In the presence of Nb33, the biased ligands were able to induce a similar effect than BU72 and DAMGO alone **(F)**. However, the presence of the two peaks with different intensity suggest substantial slow conformational exchange between different states. Of note the peak from the unliganded state (2.15 ppm) totally disappeared in the ternary complex samples for non-selective agonists (BU72, DAMGO). The NMR cross-peak of M205$^{4.63}$ in the HMQC spectra is highly dependent of the bound ligand. The distinct effects between agonists may be due their different chemotypes (**Figure 1A**). However, binding of ligands results in the appearance with two new close peaks at the downfield resonance in the $^{13}$C dimension together with a highly broadening/splitting of the apo peak (BU72), in the increase and broadening towards downfield $^{13}$C shift signal intensity (DAMGO), in the loss of signal intensity (e.g., oliceridine, buprenorphine) or no changes (e.g., PZM21) (**M**). As observed with M65$^{1.29}$, binding of Nb33 at the ICD causes more drastic changes in the NMR resonance of M205$^{4.63}$. Particularly for all agonists, we observe the appearance of multiple peaks, which shifts downfield from the apo-state peak for both $^{1}$H and $^{13}$C dimensions, recovering the new peak observed in the binary complex with BU72 (**O**). These observations suggest that the local conformation may become more dynamics and multiple conformations that exchange on the intermediate or slow NMR timescales.

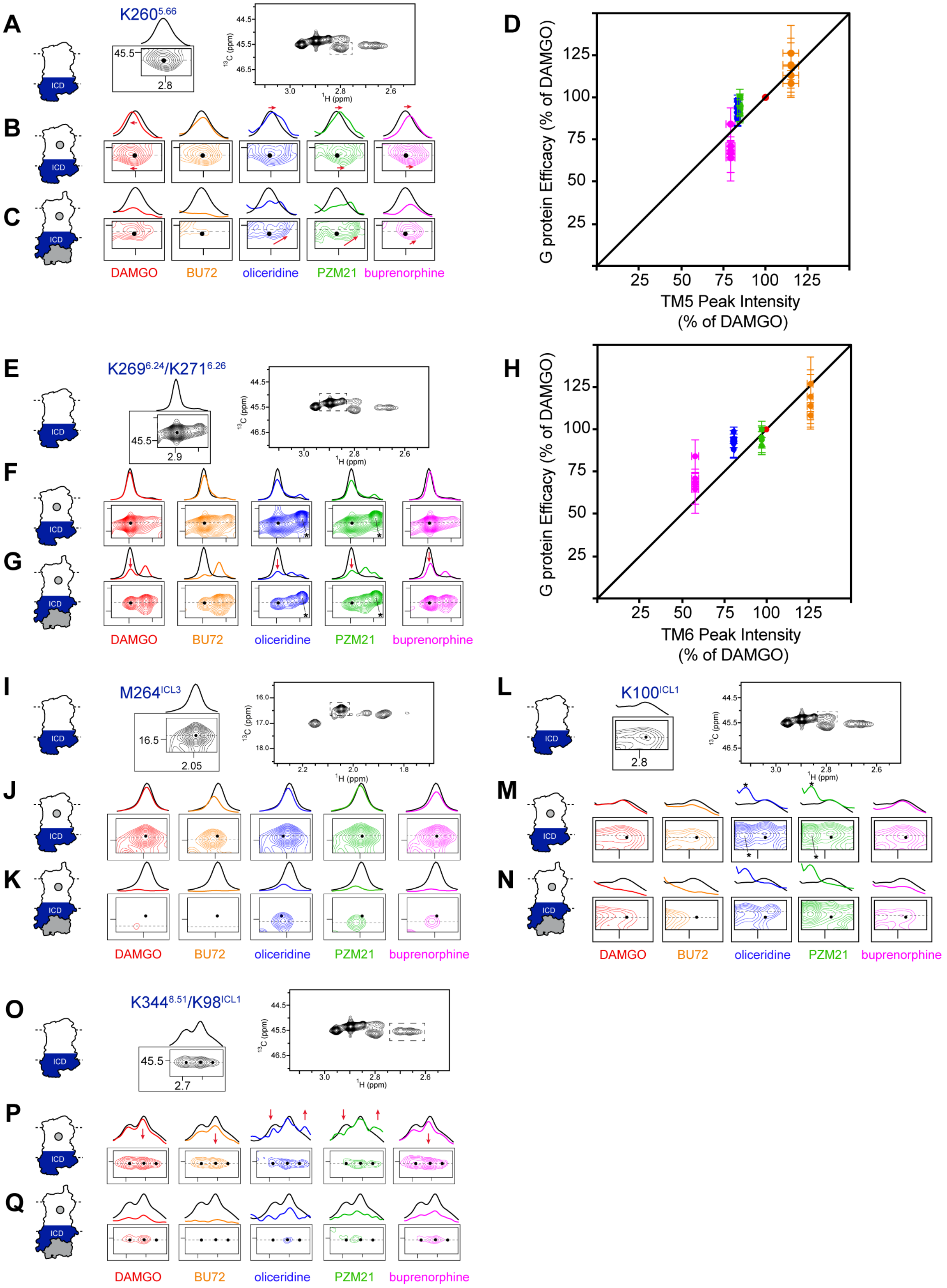

A
K260^5.66
45.5
2.8
ICD
13C (ppm)
1H (ppm)
B
C
DAMGO
BU72
oliceridine
PZM21
buprenorphine
D
G protein Efficacy (% of DAMGO)
TM5 Peak Intensity (% of DAMGO)
E
K269^6.24/K271^6.26
45.5
2.9
F
G
H
TM6 Peak Intensity (% of DAMGO)
I
M264^ICL3
16.5
2.05
J
K
L
K100^ICL1
2.8
M
N
O
K344^8.51/K98^ICL1
45.5
2.7
P
Q

**Figure S5. NMR spectral changes on the intracellular coupling domain upon binding with each agonist and Nb33 at saturating concentrations. Related to Figure 5.**

Extracted HMQC spectra of the sensors (**A—C**) K260$^{5.66}$, (**E—G**) K269$^{6.24}$/K271$^{6.26}$, (**I—K**) M264$^{ICL3}$, (**L—N**) K100$^{ICL1}$ and (**O—Q**) K344$^{8.51}$/K98$^{ICL1}$ are compared with *apo* μOR (black) (top panel), either bound to agonist (middle panel) or to agonist and Nb33 (bottom panel). Dashed black lines indicate the position of the cross-sections shown on top of the spectra. Black dots indicate the peak centers of *apo* μOR. Asterisks in (M) and (N) indicates signals from impurity most likely comes from the uncleaved N-terminus. Agonist efficacies for G proteins correlate with the changes in the peak intensities at (**D**) K260$^{5.66}$ and (**H**) K269$^{6.24}$/K271$^{6.26}$ in the ternary agonist-μOR-Nb33 complexes. DAMGO is used as reference (set at 100%). Verticals error bars indicate SEM for each of the five Gαi/o proteins (Table S1). Horizontals error bars represent the uncertainty in volume determination due to random noise.

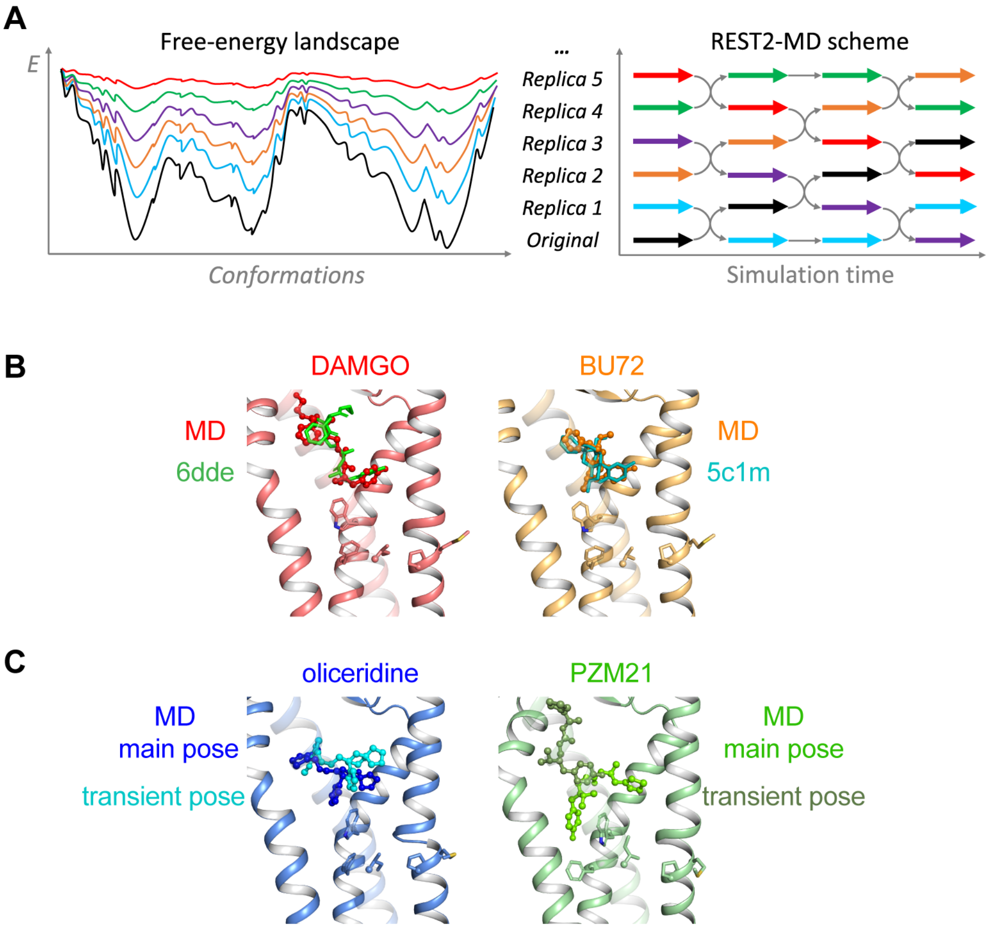


**Figure S6. REST2-MD simulation scheme and ligand binding poses captured by REST2-MD. Related to STAR Methods.**

(**A**) REST2-MD performs many replicas of the same simulation system simultaneously. The replicas have flatter free energy surfaces to ease barrier crossing. By frequently swapping the replicas during the MD, the simulations "travel" on different free energy surfaces and easily visit different conformational zones. (**B**) The binding poses of DAMGO and BU72 reproduced those in the experimental structures (PDBs 6DDE and 5C1M, respectively). (**C**) Transient binding poses of oliceridine and PZM21 compared with the main poses.

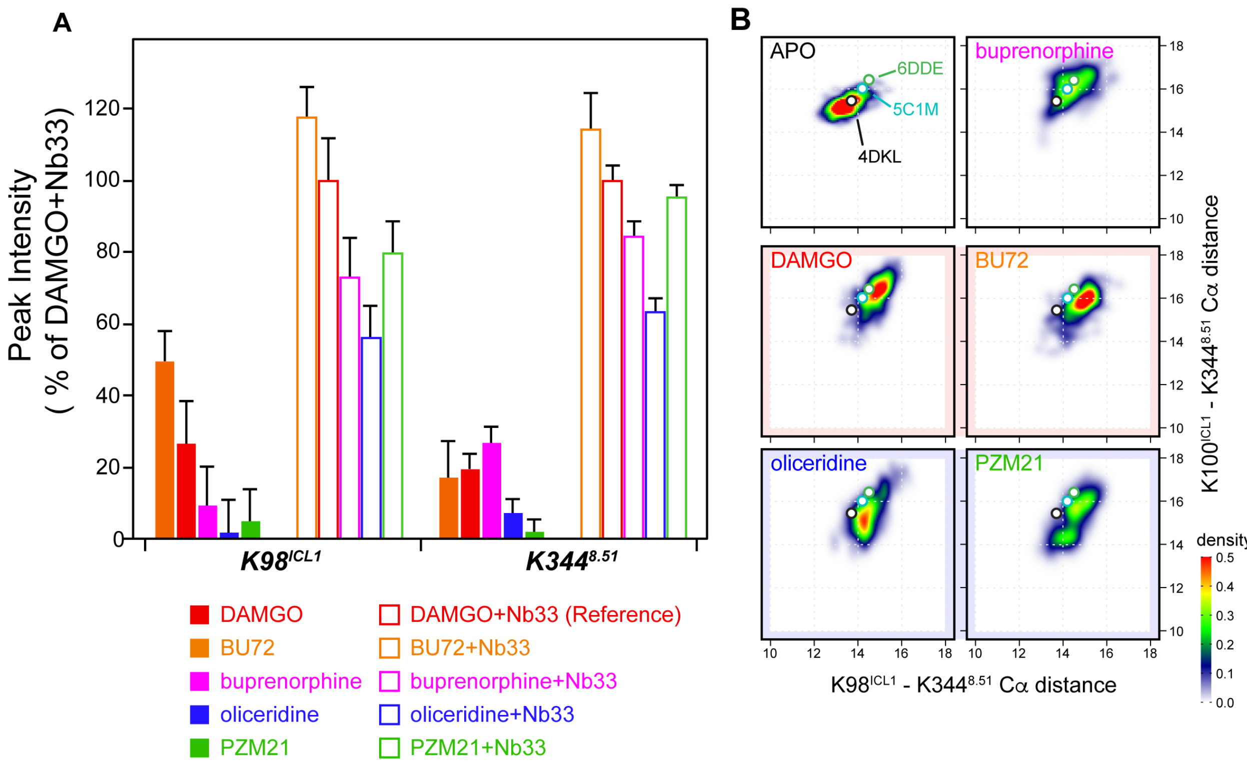


**Figure S7. Conformational changes of ICL1/H8 upon binding the agonists and Nb33. Related to Figure 5.**

(**A**) Peak intensities were normalized to the volume difference between the *apo* state and DAMGO-μOR-Nb33 ternary complex as the 100%. Error bars represent the uncertainty in volume determination due to random noise. (**B**) Density maps of the Cα distances between $K98^{ICL1}$, $K100^{ICL1}$ and $K344^{8.51}$ during REST2-MD of the agonist-μOR-Nb33 ternary complexes, compared with the *apo* state and the values in experimental structures ((PDBs 4DKL, 6DDE and 5C1M).
